# Revisiting Hydractiniidae (Hydrozoa): Mitogenomic Insights into the Diversity, Systematics and Life-Cycle Evolution of North Atlantic and Arctic Species

**DOI:** 10.64898/2026.10.09.757902

**Authors:** Lara Maleen Beckmann, Aino Hosia, Marta Ronowicz, Luis Martell, Joan J Soto-Angel

**Affiliations:** Tjärnö Marine Laboratory, University of Gothenburg, Laboratorievägen 10, 45296, Strömstad, Sweden; Gothenburg Global Biodiversity Centre, P.O. Box 463, 40530 Gothenburg, Sweden; Department of Natural History, University Museum of Bergen, P. O. Box 7800, 5020, Bergen, Norway; Marine Ecology Department, Institute of Oceanology Polish Academy of Sciences, ul. Powstancow Warszawy 55, 81-712, Sopot, Poland

**Keywords:** life-cycle strategies, integrative taxonomy, DNA barcoding, mitogenomics, developmental biology, medusa

## Abstract

Hydractiniidae is a diverse family of colonial hydrozoans exhibiting substantial morphological variation and a wide range of life-cycle strategies, including species with and without a medusa stage. Several species are established model organisms, yet fundamental gaps in current knowledge on hydractiniid biosystematics hinder comparative studies of reproductive and developmental traits and our understanding of their true distributions. Here, we employ an integrative approach combining detailed morphological assessments with DNA barcoding and complete mitochondrial genomes to reassess species diversity and distribution, phylogenetic relationships, and life-cycle variation in Hydractiniidae. Our results confirm 10 species are present in the North Atlantic and Arctic Ocean, occurring from shallow waters to 2600 m depth and from coastal fjords to deep-sea hydrothermal vents. We establish a new genus for Arctic hydractiniid species (*Bonneviactinia* gen. nov.) and reassign two species to *Schuchertinia*. We further show that some historical species records require reassessment. Mapping life cycle strategies onto the phylogeny reveals a broad pattern of increasing medusa development, consistent with multiple independent loses of the medusa stage in several clades, rather than a stepwise gain of medusa characters in Hydractiniidae. Furthermore, we clarify the identity of *Podocoryna carnea* and, through rearing experiments from polyp stage to mature medusae, characterize its morphological variation and developmental stages. By linking validated records to specimen vouchers and molecular data, we establish reference resources that will support future taxonomic, phylogenetic and biodiversity research, including molecular surveys of marine communities. Together, these results highlight the value of integrating morphological and molecular evidence, including whole mitogenomes, to resolve hydractiniid diversity and evolutionary relationships, while providing a robust comparative framework for investigating the evolution of reproductive strategies and life-cycle complexity across Medusozoa.

## 1 Introduction

Some commonly encountered and extensively studied groups can encompass surprisingly unresolved diversity. This is particularly true among many marine invertebrates, where simple morphologies, complex life histories, and historically fragmented descriptions of different life-cycle stages can obscure species boundaries, even in well-studied groups. Hydrozoans exemplify this problem, combining remarkable life-cycle diversity with often limited morphological characters for species identification. An especially striking case is the family Hydractiniidae (Agassiz, 1862), a relatively well-studied group within Hydrozoa, both widespread in marine coastal environments and widely used in biological research worldwide. In the northeastern Atlantic Ocean, hydractiniids are among the most common hydroids encountered by researchers, monitoring efforts, and the general public (Schuchert, 2008; Schuchert, 2012). In fact, species such as the “snail fur” *Hydractinia echinata* (Fleming, 1828) and the club-headed hydroid *Clava multicornis* (Forsskål, 1775) are among the few benthic hydrozoans with widely recognized common names in regional languages such as English, Swedish, and Norwegian (NBN Trust, 2026; Artsdatabanken, 2026; SLU Artdatabanken, 2026). Hydractiniid polyps grow on a wide range of substrates, from rocks, algae and gastropod shells occupied by living snails or hermit crabs (Schuchert, 2012) to artificial surfaces. Hydractiniid medusae, on the other hand, are frequent components of coastal plankton in temperate waters of the Norwegian and North Seas (Hosia and Båmstedt, 2007; Schuchert, 2012; Martell et al., 2022), as well as globally (e.g. Japan: Chaplygina & Dautova, 2005; Mediterranean: Boero & Bouillon, 1993; Gulf Stream/off Florida: Schuchert & Collins, 2024; Southwest Indian Ocean: Bourmaud et al., 2025; Argentina: Genzano et al., 2024; Greenland and Iceland: Schuchert, 2001).

Benthic hydroids are commonly associated with a wide range of organisms including bryozoans (Puce et al., 2007), kelp (González-Duarte et al., 2020), crustaceans (Widmer et al., 2010), molluscs and sponges (Puce et al., 2008) among others. These associations range from mere epibiotic to symbiotic and may be mutually beneficial. Within Hydractiniidae, colonies of *Hydractinia* spp. are said to protect their hermit-crab host (*Pagurus* spp.) against predators (Grant & Pontier, 1973), reducing interspecific competition (Fotheringham, 1976), while the polyps have an increased chance of encountering food due to the host movements (Brooks & Mariscal, 1985). Such associations may also impose costs, including reduced reproductive fitness of the host (Damiani, 2003). Furthermore, planktonic hydrozoans act as important predators in all life stages, primarily feeding on mesozooplankton (Frost et al., 2012), fish larvae and eggs, and other marine invertebrates (Purcell, 1989; Gili et al., 1998; Nicholas & Frid, 1999; Rossi et al., 2012). Fluctuating hydromedusae occurrences can also help maintain and stabilize marine ecosystem diversity (Purcell, 1989; Piraino et al., 2002), as well as contributing to benthic-pelagic coupling leading to energy transfer between ecosystems (Gili and Coma, 1998; Gili et al., 1998; Rossi et al., 2012).

In addition to their ecological significance, several hydractiniid species are well established as model systems in developmental biology and immunology research. One remarkable example is *H. echinata*, in which August Weismann (1883) found the Stammzelle, the first pluripotent cell, now known as the interstitial cell. This species remains an important model organism in studies on allorecognition and migratory stem cells (Plickert et. al, 2012). The congeneric *Hydractinia symbiolongicarpus* (Buss & Yund, 1989) is also widely used, particularly in research on gene expression and genome editing (Sanders et al., 2018). In contrast to *Hydractinia,* the closely related genus *Podocoryna* includes species with fully functional, free-swimming medusae and has therefore provided an important complementary model for studies of development, regeneration, and transdifferentiation (Galliot & Schmid, 2002; Plickert et al., 2012). However, even among model organisms, species boundaries within Hydractiniidae are not always clear-cut: for example, while the model species *Podocoryna carnea* M. Sars, 1846 has been considered to be conspecific on both sides of the Atlantic Ocean, North American populations have recently been recognized as the distinct species *P. americana* (Mayer, 1910) (Travert et al., 2023; Chang et al., 2026). Despite the attention some hydractiniid species have received in recent years, the group still suffers from poorly defined taxa, gaps in molecular data, unresolved biosystematic relationships, and frequent misidentifications of common species, often resulting from incongruences in public DNA databases. Insufficient species descriptions, similar morphology with lacking informative traits, and strong phenotypic plasticity complicate hydractiniid identification and classification (Schuchert, 2008; Miglietta et al., 2009). The latest attempts to address these challenges have relied on molecular approaches (Miglietta et al., 2009; 2010; Miglietta & Cunningham, 2012); however, for most described species, genetic data are either lacking or geographically limited, making it difficult to assess variation and assign species confidently. Miglietta et al. (2009, 2010; Miglietta & Cunningham, 2012) validated nine genera within the Hydractiniidae (*Clavactinia* Thornely, 1904; *Janaria* Stechow, 1921b; *Hydrissa* Stechow, 1921a; *Hydractinia* Van Beneden, 1844; *Schuchertinia* Miglietta, McNally and Cunningham, 2010; *Clava* Gmelin, 1788; *Stylactis* Allman, 1864, *Bouillonactinia* Miglietta, McNally and Cunningham, 2010 and *Podocoryna* M. Sars, 1846) using a combination of short DNA sequences and morphological characters, but they also emphasized the need for more comprehensive molecular datasets to clarify relationships among genera, assess life cycle variation and confidently assign species to the proposed established genus-level framework. Resolving these issues is essential for the reliable inclusion of hydractiniids in biodiversity monitoring programs, and it is also critical to ensure that research on hydractiniid model species is accurately linked to the correct taxon, allowing results to be compared and generalized across studies.

The northeastern Atlantic is particularly important for this purpose because it includes the type localities of several well-known hydractiniids, including *H. echinata, C. multicornis,* and *P. carnea* (Forsskål, 1775; Fleming, 1828; M. Sars, 1846, respectively). As type taxa for their respective genera, these species are central reference points for hydractiniid taxonomy. The region also has a long history of hydractiniid research, beginning with major nineteenth- and early twentieth-century oceanographic expeditions. The Norwegian North-Atlantic Expedition (1876-1878) provided material for the descriptions of several northern hydrozoans, including *Hydractinia minuta* Bonnevie, 1898 (now regarded as *H. carica*). The Danish Ingolf Expedition (1895-1896) and later the Godthaab Expedition (1928) generated extensive material from the deep northern Atlantic and Greenland waters, including the material from which *Hydractinia ingolfi* Kramp, 1932 was described. These and later surveys of Norwegian waters generated numerous additional records, with historical records of *H. echinata* concentrated along the southern and western coasts of Norway (Bonnevie, 1901; Broch, 1910, 1911; Christiansen, 1972; Stepanjants, 1989). However, many historical records are based on single observations or on species described from very limited material, making their identities and distributions difficult to assess. Such records were sometimes extrapolated to other geographic areas, while easily recognized species continued to dominate field identifications. This combination of incomplete historical sampling, uncertain species identifications, and uneven recognition of less conspicuous taxa may have contributed to an incomplete picture of hydractiniid diversity in the region. Public biodiversity databases retain some of these historical patterns, with observations concentrated heavily on *C. multicornis* and *H. echinata* despite the broader hydractiniid diversity reported from Norwegian waters (GBIF; Artsdatabanken, 2026). A molecularly informed inventory is therefore needed to reassess the diversity and distribution of Hydractiniidae in the northeastern Atlantic.

In this study, we provide an integrative assessment of the family Hydractiniidae in northeastern Atlantic and Arctic waters, including both coastal and deep-sea environments. Using a combination of morphological examination, multilocus molecular phylogenetics, and mitogenomics, we (1) clarify the taxonomic identities of the region’s hydractiniids; (2) compile an updated inventory with ecological and distributional information; (3) discuss new insights into life-cycle evolution in the family and (4) provide complete mitochondrial genomes and nuclear 18S and 28S sequences for multiple hydractiniid species, establishing a reference resource for future biodiversity and evolutionary studies.

## 2 Material & Methods

### 2.1 Sample acquisition

The specimens analyzed in this study originated from various sources (Supplementary Table 1), including the Invertebrate Collections of the University Museum of Bergen (UMB) and the Institute of Oceanology of the Polish Academy of Sciences (IOPAN). Fresh specimens were collected from marine localities along the Norwegian coast during projects HYPNO, NorHydro and NOAH (2015-2026), funded by the Norwegian Biodiversity Information Centre (Supplementary Table 1). The study area covers the temperate northeastern Atlantic and the European sector of the Arctic Ocean, ranging from the English Channel in the south to the waters around Svalbard in the north (Fig. 1). For comparative purposes, one specimen of *Podocoryna exigua* (Haeckel, 1880) collected from the Mediterranean Sea was also included. The material included both medusa and polyp stages from diverse environments, collected using methods such as dredging with an Agassiz trawl or triangular dredge, hand-picking (e.g., by scuba divers), and plankton sampling with various nets. Live specimens were documented through photography and videography prior to fixation in 96% ethanol, with physical and/or digital vouchers deposited at UMB. Morphological identifications were based on a combination of careful examination of the original species descriptions, the summary accounts by Schuchert (2008, 2012), and the study of museum material. This included specimens and type material of the species *Bonneviactinia allmanii* (Bonnevie, 1898) gen. nov., comb. nov. (ZMO_B599 and ZMO_B600) and *Schuchertinia sarsii* (Steenstrup, 1850) comb. nov. (ZMO_B686 and ZMO_1347). Freshly collected, reproductive polyp colonies of *Podocoryna carnea* and *Podocoryna areolata* from western Norway, near the type locality of the species, were cultured in the invertebrate laboratory at UMB. The colonies were held in seawater tanks for one night after collection until medusae were released. The released medusae were then kept in 6 L containers with a gear motor connected to serological pipette rotating at 15 rpm as described in Soto-Angel et al. (2024). Animals were kept at approximately 12°C and fed with *Brachionus* and *Artemia* every other day. Medusae development was monitored daily and photographed every other day. Both cultured strains (*P. areolata* and *P. carnea*) were sequenced for 16S and COI (Sanger sequencing), and genomic data was obtained to reconstruct the mitogenome (see below).

**Figure 1.**
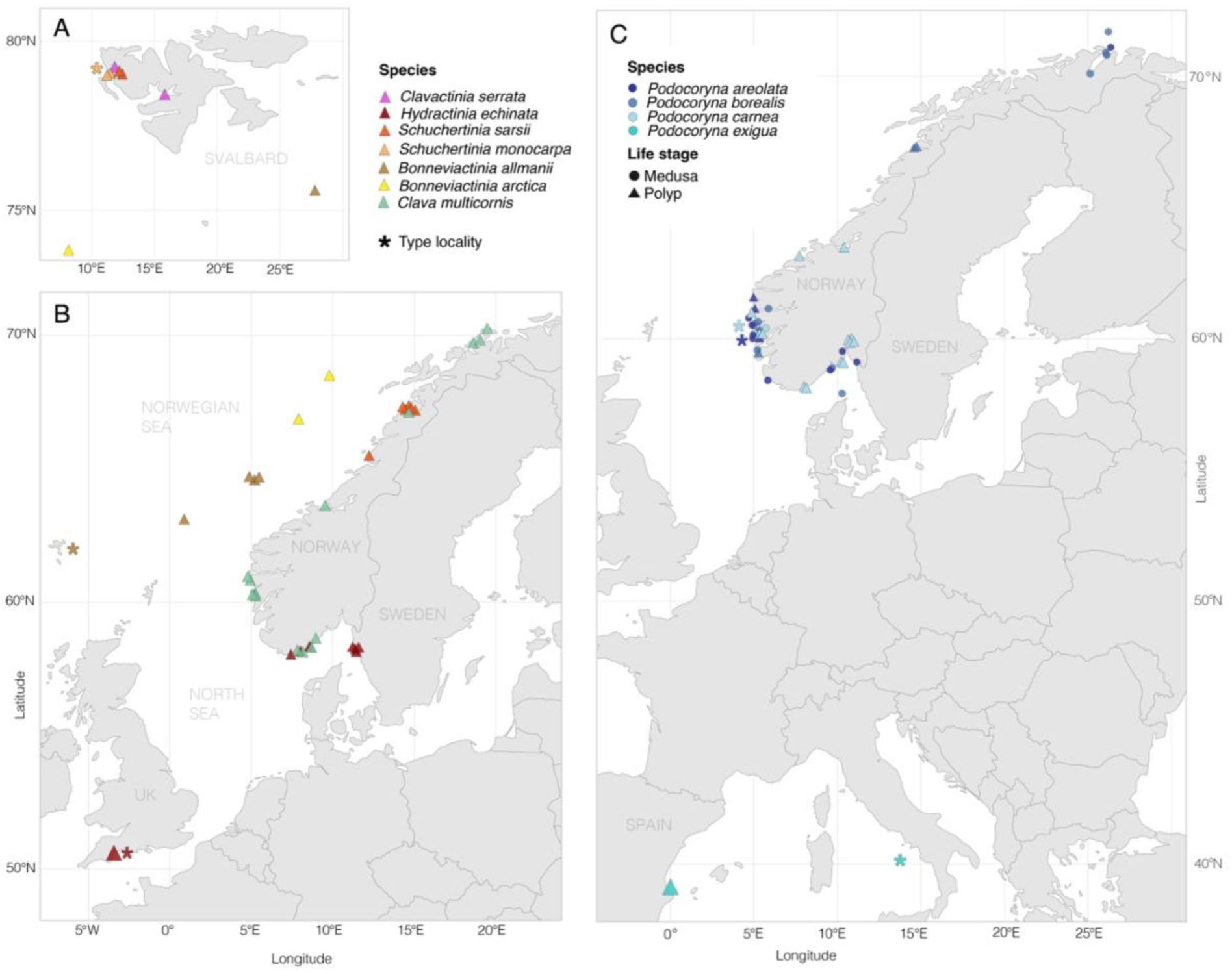
Geographic distribution of examined hydractiniid specimens. Species are indicated by colour, and the life stage at collection by shape (medusa = circle; polyp = triangle). Type localities are indicated by an asterisk (*) in the corresponding species colour where they fall within the mapped area. (A) Hydractiniid specimens collected around Svalbard. (B) All non-*Podocoryna* specimens collected. (C) *Podocoryna* specimens collected.

### 2.2 DNA extraction, amplification, and sequencing

A subset of 82 specimens (Supplementary Table 1) was selected for molecular characterization, with the aim of including all taxa and achieving the widest possible geographic and temporal coverage. For each of these specimens, a small tissue sample was dissected: for medusa stages, the umbrella margin was used, while for polyp stages, either tentacles or entire polyps were sampled. DNA extractions were then performed at either the Canadian Centre for DNA Barcoding (CCDB), following the protocols of Ratnasingham and Hebert (2007), or at the University of Bergen DNA laboratory, where samples were processed for either DNA barcoding or Whole Genome Sequencing (WGS) and mitogenome reconstruction. Details on the pipeline chosen for each specimen are provided in Table S1 in the supplementary material.

Genomic DNA (gDNA) was extracted for the specimens selected for DNA barcoding using the QuickExtract™ DNA Extraction Solution Kit, following a modified protocol after Nygren et al. (2018). Three genetic markers (mitochondrial COI and 16S, nuclear ITS) were amplified under the PCR conditions detailed in Table 1 and as described in Beckmann et al. (2023). PCR products were assessed by electrophoresis on 1% agarose gels, purified with ExoSAP-IT. Whenever COI amplifications yielded multiple bands, the target product (∼650 bp) was excised from the agarose gel using the QIAGEN MinElute Gel Extraction Kit. The resulting PCR products were sequenced at the University of Bergen sequencing facility or through Azenta Life Sciences. For all obtained sequences, chromatograms were analyzed in FinchTV, compared against the NCBI database to detect contamination, and manually edited to solve ambiguities. Consensus sequences were assembled in Geneious v.11.1.5.

**Table 1.**
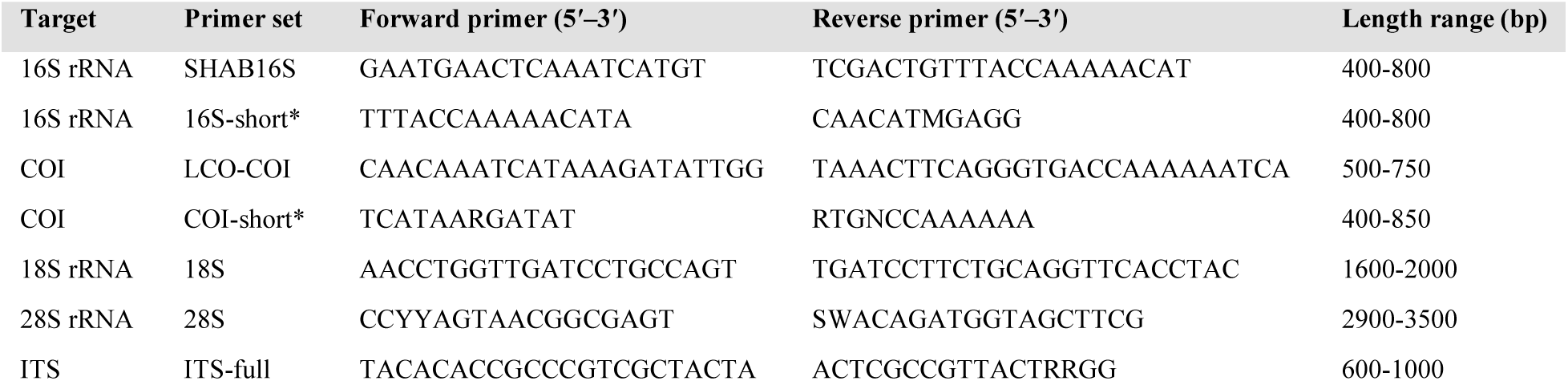
Primer pairs used for *in silico* PCR (sPCR) with Sharkmer to extract mitochondrial and nuclear gene regions from WGS data. Minimum and maximum expected amplicon lengths used for sPCR are indicated. Shortened primer pairs used when the corresponding full-length primer pair did not result in detection of the target gene are indicated by an asterisk (*).

Specimens selected for WGS, were extracted using the DNeasy Blood and Tissue kit (QIAGEN) following the manufacturer’s instructions. For those challenging samples with very low DNA yield due to unknown/complicated preservation history, single-stranded library preparations were prepared using SRSLY® PicoPlus Uracil+ Base Kit from Claret Bioscience (cat. no. CBS-K250B-U-24), a ligation-based library preparation method. In brief: Libraries were constructed following the manufacturer’s protocol. Total input of gDNA varied between samples, from unquantifiable to 15 ng. DNA was heat-denatured and adapters were ligated to the single strands (Troll et al. 2019). Ligation products were purified and size-selected using 59.4 µl of Sera-Mag Select beads (Cytiva), aiming for small fragment purification. Libraries were then amplified through index PCR with 8-13 cycles, depending on the input amount following the protocol’s recommendations. Unique dual (UDI) index primers were supplied by Claret Bioscience. A second bead cleanup using 50 µl of Sera-Mag Select beads followed, aiming for small fragment retention, and using single-sided magnetic bead purification protocol for indexed libraries with less than 10 ng of Template DNA Input.

### 2.3 Whole Genome Sequencing data processing: sPCR, mitogenome assembly and annotation

To obtain mitochondrial genomes and additional nuclear markers (18S, 28S), a subset of samples was subjected to low-coverage WGS aiming at 5-10x coverage. Sequencing libraries were prepared by Novogene following standard Illumina protocols and sequenced on an Illumina NovaSeq X Plus platform, generating paired-end 150-bp reads (PE150).

Raw reads were quality-controlled using fastQC (Andrews, 2010) and adapter-trimmed using fastp (Chen, 2025). Reads shorter than 25 bp after trimming were discarded (-l 25) and paired-end adapter detection was enabled (--detect_adapter_for_pe). Quality filtering was otherwise performed using the default fastp settings. Quality-control reports generated by fastQC were inspected following filtering.

To extract individual mitochondrial and nuclear genes from the WGS data, *in silico* PCR (sPCR) was performed using Sharkmer (Dunn & Church, 2026). Standard cnidarian primer pairs, as well as modified hydrozoan-specific primer pairs, were used to target 16S rRNA, cytochrome c oxidase subunit I (COI), 18S rRNA, 28S rRNA, and the internal transcribed spacer (ITS) regions (Table 1). For 16S and COI, both full-length and shortened primer variants were used to improve recovery of target sequences from individual samples. Up to 4 million reads per sample were screened for target sequences.

Mitochondrial genomes were assembled *de novo* using GetOrganelle v1.7.7.1 (Jin & Yu et al, 2020), using the animal_mt workflow. Assemblies were performed with a maximum of 15 extension rounds (-R 15), no hard upper limit on the number of reads (--max-reads inf), k-mer sizes of 21, 45, 65, 85, and 105 (-k 21,45,65,85,105), and a word size of 55 (-w 55). A reference mitochondrial sequence was provided as the initial seed (-s) (*Hydractinia echinata*; Accession number: CM057788; Schnitzler et al., 2024). Putative mitochondrial contigs were identified based on similarity to reference Hydractiniidae mitogenomes available on NCBI. Annotation of mitochondrial genomes was performed using MITOS2 (Bernt et al., 2013). Predicted gene boundaries and annotations were manually curated, compared/aligned with published anthoathecate mitogenomes and checked for STOP and START codons. Transfer RNA genes were independently evaluated using tRNAscan-SE 2.0 (Lowe & Chan, 2016). Gene boundaries, orientation, and order were verified through comparison with published anthoathecate mitogenomes of *H. echinata* (CM057788), *H. symbiolongicarpus* (LN901197), *H. polyclina* (LN901196), *P. americana* (LN901210), and *C. multicornis* (JN700935) (details in Supplementary Table 2).

### 2.4 Sequence dataset and alignment

Reference sequences for 16S, 18S, and 28S rRNA genes from Hydractiniidae were retrieved from the NCBI database for comparative analyses (Supplementary Table 2). Two datasets were constructed: (1) a comprehensive 16S dataset including global Hydractiniidae diversity (270 sequences), and (2) a reduced dataset including 1-3 selected sequences of each of the markers per phylogenetic clade identified in (1).

The concatenated dataset used for genus-level analyses included all available genera represented by 16S, 18S, and 28S markers (Supplementary Table 2). The data came from a single individual, except for *Bouillonactinia* spp., where complete marker coverage was not available. Here, chimeric concatenations were constructed from different specimens of the same species (Supplementary Figure 2). These chimeric sequences were used solely to assess the potential phylogenetic placement of the genus and were not included in the final hypothesis testing. Stylasteridae was selected as outgroup in all analyses, based on the phylogenetic proximity of this family to Hydractiniidae (Pica et al, 2025). All available hydractiniid mitochondrial genomes were also retrieved from NCBI for comparative analyses (Kayal et al, 2012; 2015). Sequence alignments for each marker were generated using MAFFT as implemented in Unipro UGENE v53.1. Alignments were visually inspected and manually adjusted where necessary (trimmed at START/STOP codons). All newly generated sequences have been deposited in BOLD and GenBank (accession numbers provided in Supplementary Table 1).

### 2.5 Phylogenetic analysis

Phylogenetic relationships were inferred using Maximum Likelihood (ML) as implemented in IQ-TREE v2 (Nguyen et al., 2015; Trifinopoulos et al., 2016). For concatenated datasets, partitions were defined by gene, and the best-fitting substitution models were selected using ModelFinder as implemented in IQ-TREE. Where applicable, partition merging was performed using the MFP+MERGE option. Branch support was assessed using ultrafast bootstrap approximation (UFBoot) with 1000 replicates (Hoang et al., 2017) and SH-like approximate likelihood ratio tests (SH-aLRT) with 1000 replicates. Analyses were run with four discrete gamma rate categories. Separate phylogenetic analyses were conducted for (1) the 16S dataset including global Hydractiniidae diversity, (2) the concatenated nuclear and mitochondrial marker dataset, and (3) mitochondrial genome sequences. For 3) all available genes were isolated from the mitogenome and then concatenated. All resulting trees were visualized and edited in FigTree v1.4.4. Final figures were prepared in Adobe Illustrator v25.4, with adjustments limited to text, coloration, and layout; tree topology and branch lengths were not altered.

### 2.6 Morphological characterization and host identification

A set of diagnostic characters with potential for species delimitation were evaluated in the collected specimens. These characters were selected based on previous taxonomic analysis in Hydractiniidae due to their potential for the morphological characterization of the identified clades (Schuchert, 2008, 2012; Miglietta et al., 2010; Miglietta & Cunningham, 2012). For each specimen, the corresponding state for each character was recorded and measured when appropriate (with ImageJ; Schneider et al., 2012). For the polyp stage, these characters included polyp size, tentacle number, tentacle arrangement, shell size of the host, and spine morphology and size. Depending on the density of the colony 1-5 polyps (gastrozooids) were measured per sample. The size of each polyp was measured from the mouth to the aboral end without tentacles. Note that the measurements were conducted in preserved specimens (ethanol). The host or substrate of the colonies were identified to species level when possible. The spine size was either measured under the microscope or estimated with the aid of the tip of a fine lab pincer (ESD-11 pincer has a tip size of 0.2 mm) when removal was not possible. The number of tentacles was counted and averaged for five mature gastrozooids per colony.

The evaluated characters for the medusae stage were umbrella height, tentacle/tentacular bulb number, tentacle arrangement, the position of gonads, size of gonads, shape of the manubrium, and general shape of the umbrella. Umbrella height was measured from the aboral end to the oral end, and the tentacles were counted when they were visible at the tentacle bulbs.

## 3 Results

### 3.1 Phylogenetic analysis

The 16S phylogeny recovered 50 well-supported and distinct clades corresponding to putative species-level hydractiniids, including all the sequences produced in this study and those mined from NCBI (Fig. 2; Supplementary File 1 for all available hydractiniid sequences). Of those, ten species-level clades were represented by material originating from the study area: *Hydractinia echinata, Clava multicornis, Podocoryna areolata, Podocoryna borealis, Podocoryna carnea, Schuchertinia sarsii* comb. nov.*, Schuchertinia monocarpa* comb. nov*., Clavactinia serrata, Bonneviactinia arctica* gen. nov., comb nov., and *Bonneviactinia allmanii* gen. nov., comb. nov. Species assignments were based on congruence across markers and datasets. The diagnoses of the new genus and the rationale for the new combinations are provided in the discussion.

**Figure 2.**
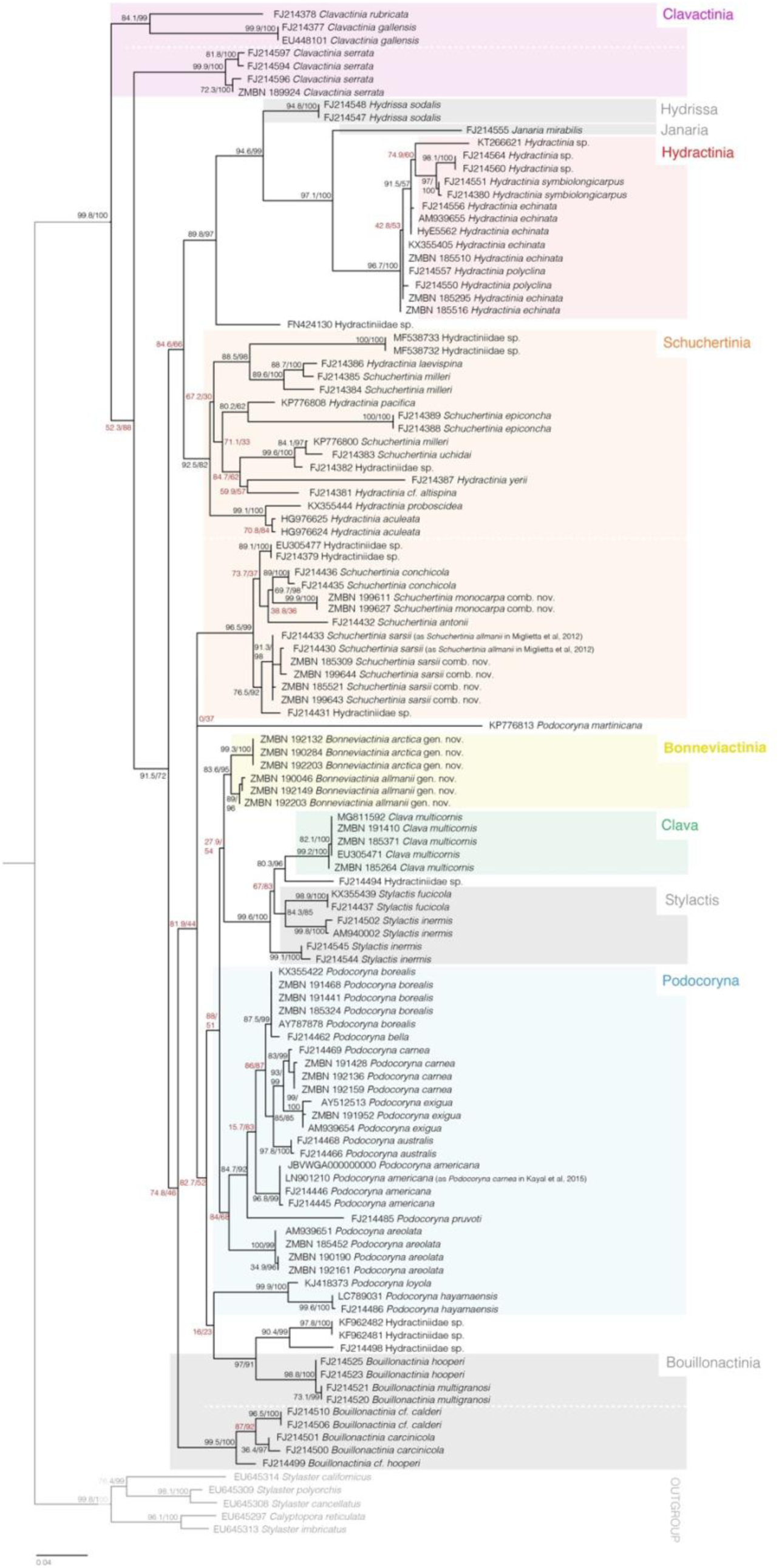
Maximum-likelihood phylogeny inferred from the 16S dataset using IQ-TREE and using a reduced dataset with a few specimens per clade. Node labels indicate ultrafast bootstrap support followed by SH-aLRT support (UFBoot/SH-aLRT).

At the global scale, three species-level clades were recovered within *Clavactinia*: *C. rubricata, C. gallensis,* and *C. serrata*. The genus was not resolved as monophyletic, with *C. serrata* forming a separate lineage from *C. rubricata* and *C. gallensis* (Fig. 2, Supplementary File 1), with high support for both major clades (84 UFBoot / 99 SH-aLRT and 99/100, respectively). Within *C. serrata,* two distinct clades were recovered. One included the sequence generated in this study from Svalbard and a sequence from the Bering Sea, USA. The second comprised sequences from the Bering Sea and Washington State. These specimens were collected from material growing on hermit crabs rather than molluscs.

The genera *Hydrissa* and *Janaria* were recovered with high support (94/100 and 97/100, respectively), including *H. sodalis* and *J. mirabilis*. The genus *Hydractinia* included four to five putative species-level clades, including at least two species-level clades without current species-level identifications (*Hydractinia sp.* indet.). *Hydractinia symbiolongicarpus* was recovered with high support (97/100), and the nearest neighbor being an undescribed *Hydractinia* sp., and clearly separated from *H. echinata*. *Hydractinia echinata* was recovered as a species complex including sequences from the type locality in the UK, Norway and the USA. The sequences grouped in two closely related clades, with US localities grouped together but closely related to the European ones, showing little divergence. The *H. echinata* clade included sequences identified as *H. polyclina* from France and the USA.

The genus *Schuchertinia* was resolved as paraphyletic in the 16S-phylogeny (Fig. 2; Supplementary File 1) and with long-branch in the concatenated analysis (Fig. 3). We further refer to the two resolved clades as *Schuchertinia* clade I and II. *Schuchertinia* clade I includes the type species *S. epiconcha* with support of 92/82, as well as *S. laevispina, S. milleri*, *S. uchidai, S. proboscidea*, and *H. aculeata*. Sequences identified as *S. milleri* occurred in three distinct clades, suggesting either misidentification of some specimens or species-level diversity within the name *S. milleri. Schuchertinia* clade II was strongly supported (96/99) and included the Norwegian species recovered in this study, *S. sarsii* comb. nov. and *S. monocarpa* comb. nov, as well as *S. antonii, S. conchicola* and two undetermined Hydractiniidae. Within clade II, the *S. sarsii* material identified in this study with reference to type material grouped with GenBank sequences labelled as *S. allmanii*, including material from Iceland and the Bering Sea. This placement indicated a likely misidentification of two specimens originally assigned to *S. allmanii* by Miglietta & Cunningham (2012), which instead fall within *S. sarsii* comb. nov. Sequences obtained from specimens morphologically identified as *Hydractinia monocarpa* were also placed within the *Schuchertinia* clade II lineage and were therefore assigned to *Schuchertinia monocarpa* comb. nov. Mitogenome and slow-evolving nuclear marker data support this (see below).

**Figure 3.**
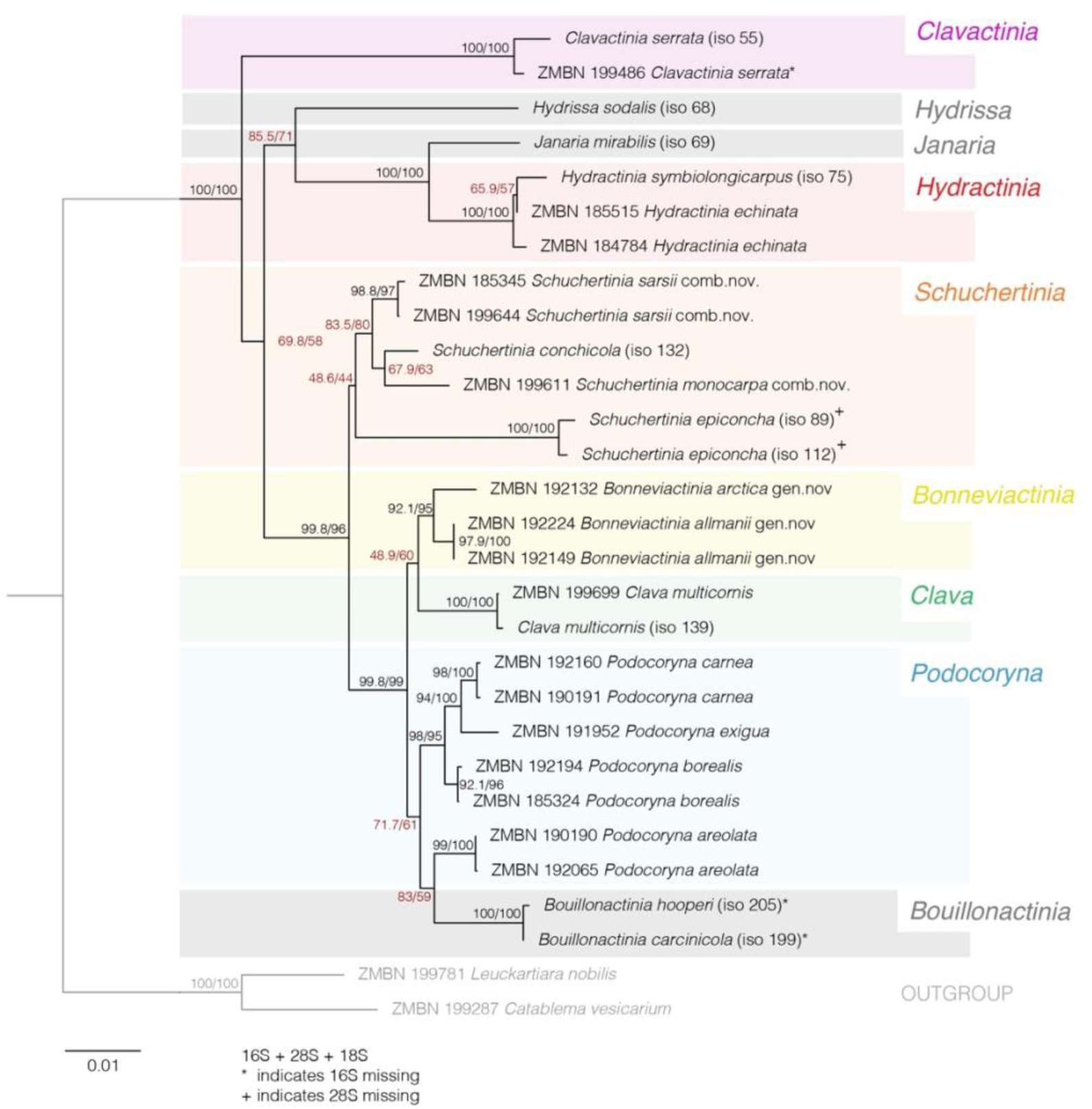
Maximum-likelihood phylogeny inferred from the concatenated 16S, 28S and 18S dataset using IQ-TREE. Node labels indicate ultrafast bootstrap support followed by SH-aLRT support (UFBoot/SH-aLRT). The asterix (*) indicates missing 16S; + indicates missing 28S.

In the concatenated phylogeny including 16S, 18S, and 28S data (Fig. 3), *S. epiconcha* from clade I remained clearly separated from clade II. However, support for the placement of the *S. epiconcha* lineage within *Schuchertinia* was low (48/44; Fig. 3), and the two specimens lacked full-length 28S sequences. Thus, the current multilocus data did not resolve the genus-level limits of *Schuchertinia*. Because sufficient data were not available to disentangle the genus, and the type material of *S. epiconcha* was not examined, no further changes to the genus are introduced in this study. The phylogenies nevertheless recovered several putative species-level lineages within *Schuchertinia*, including several lineages without current species-level identifications.

*Clava* was recovered as a well-supported monophyletic genus represented by the single species *Clava multicornis* (99/100). Sequences from France, the USA, Canada, Iceland, and Norway were included in the same *C. multicornis* clade, with little genetic difference observable in the tree. *Stylactis* was recovered with three species-level clades. Two of these contained sequences identified as *S. inermis*, indicating either potential misidentification or the presence of two distinct species currently sharing this name. One unidentified species-level lineage was placed between *Clava* and *Stylactis.* No representatives of *Stylactis* were identified from the study region.

*Podocoryna*, the medusa-bearing genus in the family, included nine putative species-level clades in the 16S dataset but was not recovered as monophyletic in either the 16S or concatenated phylogeny (Figs. 2 and 3). In the 16S phylogeny, *Podocoryna loyola* and *Podocoryna hayamaensis* were clearly distinct from the other *Podocoryna* lineages. In the concatenated phylogeny (Fig. 3), which did not include *P. loyola* and *P. hayamaensis, P. areolata* did not fall within the rest of the *Podocoryna* species but instead grouped with *Bouillonactinia*, which had incomplete data (missing 16S). Within the *Podocoryna* clade, *P. borealis* formed a distinct lineage containing sequences from Iceland, Scotland, Norway, and New Zealand. The New Zealand sequence had previously been identified as *Podocoryna bella*. 16S sequences from Iceland, one from Scotland, one from Norway, and the New Zealand sequence clustered within this lineage. Several sequences obtained from polyp material were also assigned to the same genetic clade as the sequenced *P. borealis* medusae, providing the first polyp sequence/barcode for this species in Norway. *Podocoryna carnea* was clearly distinct from *P. exigua* and *P. americana*. Model-organism sequences JBVWGA000000000 (Chang et al., 2026) and LN901210 (Kayal et al. (2015) clustered with *P. americana.* Two specimens assigned to *P. americana* were also identified as likely misidentifications, having previously been assigned to *P. carnea* by Miglietta et al. (2009) and Kayal et al. (2015). *Podocoryna areolata* formed a monophyletic clade with little intraspecific divergence. Only one additional 16S sequence from Norway was available. Several sequences from previously unidentified polyp samples were assigned to *P. areolata.* The species is included for the first time in a broader phylogenetic analysis, where it consistently formed a distinct lineage from the other *Podocoryna* species recovered here.

The genus *Bouillonactinia* was not monophyletic in the 16S phylogeny, with two distinct, well supported clades. One clade included *Bouillonactinia hooperi* and *Bouillonactinia multigranosi*, whereas the second included *Bouillonactinia cf. calderi, Bouillonactinia carcinicola,* and *Bouillonactinia cf. hooperi.* The latter *B. cf. hooperi* was not closely related to *B. hooperi* in clade I. In the concatenated phylogeny (Fig. 3), *Bouillonactinia* appeared monophyletic with *Podocoryna*, but with low support in both the concatenated analysis and the chimera analysis (Supplementary File 2). The two species that could be included in the concatenated analysis were representatives of the two resolved *Bouillonactinia* clades mentioned above (*B. hooperi* and *B. carcinocola*). However, those two were represented by incomplete data, and thus the placement of *Bouillonactinia* was unstable (no 16S). Analyses based on 18S and 28S suggested an affinity with *Podocoryna,* whereas inclusion of chimeric sequences placed it outside this clade, again with low support (Supplementary File 2). No species of *Bouillonactinia* were found in the study area.

A distinct lineage containing the material assigned to *Schuchertinia allmanii* and *Hydractinia arctica* was recovered in all trees (Figs. 2-4, Supplementary File 1 and 2). The lineage was supported by >80% bootstrap support in the 16S phylogeny (Fig. 2), >95% in the concatenated phylogeny (Fig. 3), and 100% in the mitogenome phylogeny (Fig. 4). The lineage was most closely related to *Clava*. The two species are therefore transferred to *Bonneviactinia* gen. nov., resulting in the combinations *Bonneviactinia allmanii* gen. nov., comb. nov. and *Bonneviactinia arctica* gen. nov., comb. nov. No additional specimens nor sequences publicly available were identified as belonging to this newly resolved lineage.

**Figure 4.**
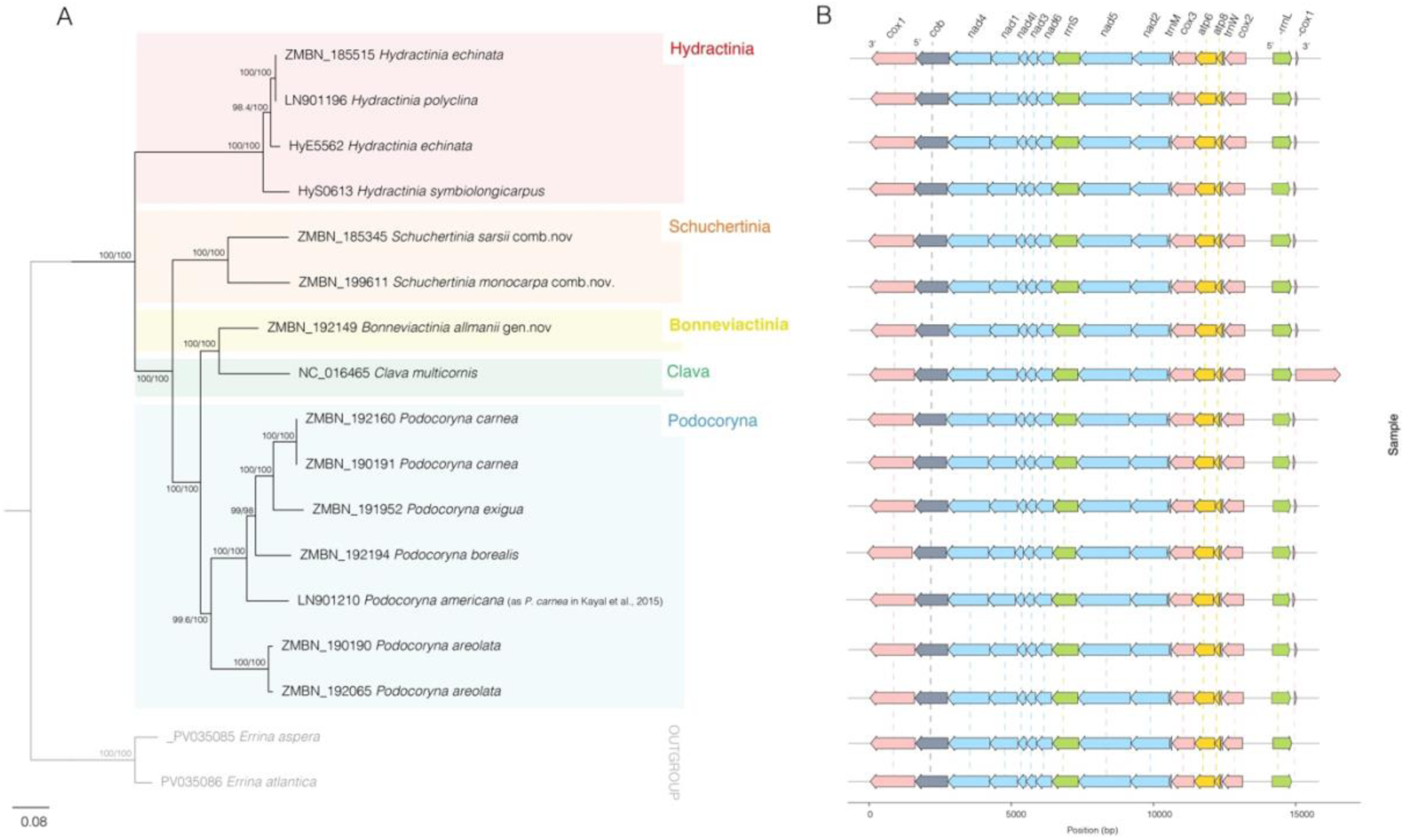
A) Maximum-likelihood phylogeny inferred from the concatenated mitogenome-level dataset using IQ-TREE. Node labels indicate ultrafast bootstrap support followed by SH-aLRT support (UFBoot/SH-aLRT). B) Mitochondrial gene orientation and organisation of the specimen used in the phylogenetic tree.

The relationships within several genera, particularly *Schuchertinia, Podocoryna,* and *Bouillonactinia*, remained unresolved due to incomplete data and/or taxon sampling. But overall, the concatenated 16S, 18S, and 28S phylogeny broadly supported the major relationships recovered in the 16S analysis (Fig. 3).

### 3.2 Mitogenome phylogeny and structure

Seven new mitogenomes were generated for hydractiniids: *S*. *sarsii* comb. nov*., S*. *monocarpa* comb. nov.*, P. areolata, P. borealis, P. carnea, P. exigua,* and *B*. *allmanii* comb. nov., gen. nov.

The mitochondrial phylogeny was largely congruent with the concatenated nuclear and mitochondrial marker dataset and recovered similar relationships with strong support (>99% bootstrap; Fig. 4A). The distinct lineage containing *Bonneviactinia allmanii* gen. nov., comb. nov. was recovered with 100% bootstrap support in the mitogenome phylogeny, with *Clava multicornis* as its sister taxon, corroborating its placement in the 16S and concatenated phylogenies.

Gene order and orientation were broadly consistent with the ancestral hydrozoan mitogenome arrangement described by Katal et al. (2015) (cox1, cob, nad4, nad1, nad4L, nad3, nad6, rrnS, nad5, nad2, trnM, cox3, atp6, atp8, trnW, cox2, -rrnL and -cox1) (Fig. 4B), as well as with gene arrangements reported for Filifera I and II (Ling et al., 2023) and Stylasteriidae (Pica et al., 2025). A cox1 pseudogene at the terminal end of the mitogenome was present in all specimens generated in this study and was of similar length among specimens. The *Clava multicornis* mitogenome mined from NCBI (JN700935) contained a complete duplicate of the functional cox1 gene at the “terminal end” of the mitogenome (Fig. 4B).

### 3.3 Morphological characteristics

The hydractiniids examined in this study showed variation in colony growth form, hydrorhiza, reproductive structures, spine morphology, gastropolyp size, and tentacle number (Table 1; Fig. 5-7). Species-level characteristics are summarised in Table 2, with new measurements from this study and previously published data (taken from Schuchert, 2008 and references therein).

**Figure 5.**
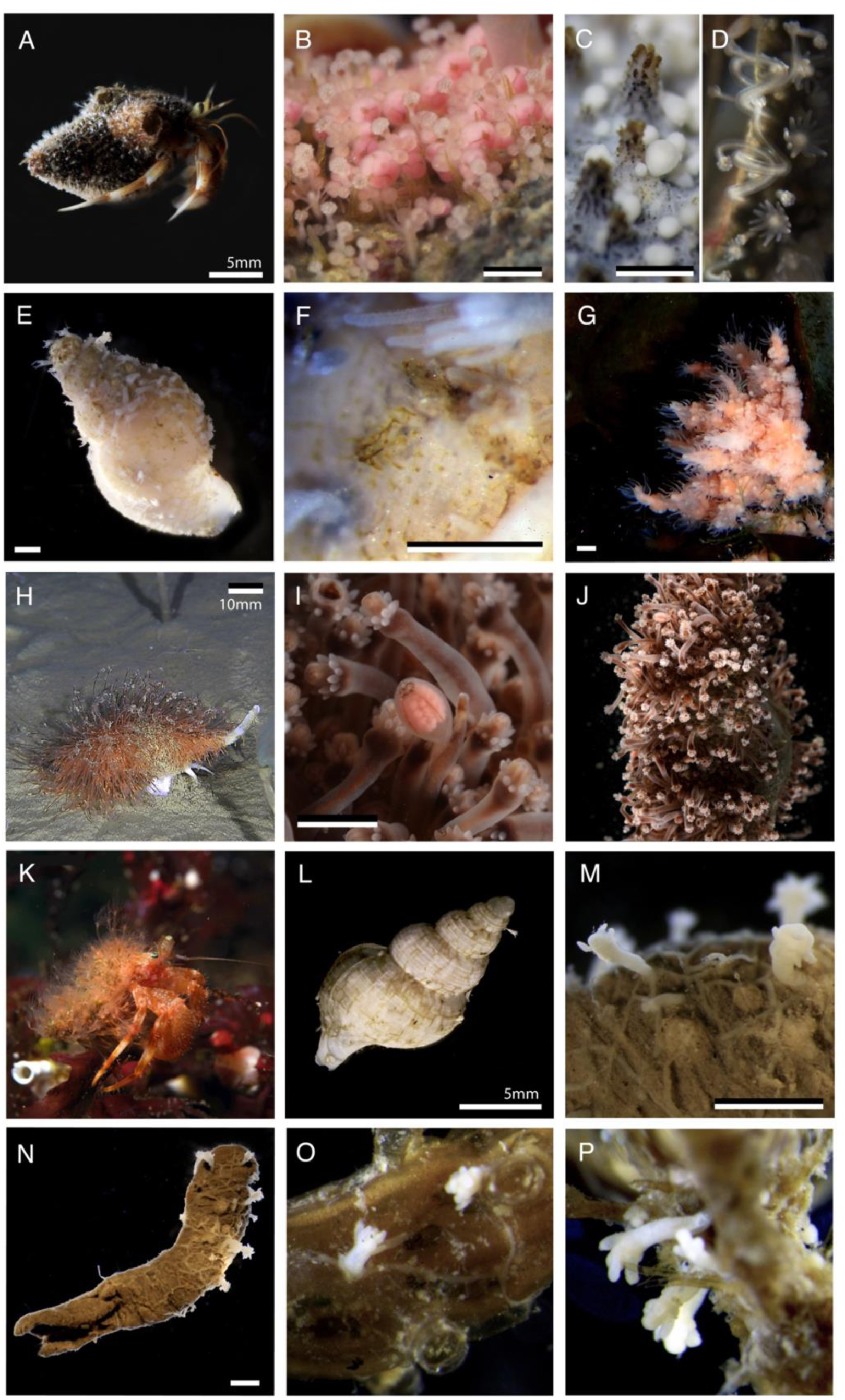
Morphology of non-medusa-bearing hydractiniid species observed in this study. (A-C) *Hydractinia echinata*. (A) Polyp colony growing on its typical host, *Pagurus bernhardus.* (B) Close-up of gastro-and gonozooids. (C) Spines showing the characteristic serration; (D) close-up of the shell rim showing dactylozooids and gastrozooids. (E-F) *Clavactinia serrata.* (E) Shell bearing a polyp colony. (F) Close-up of the short, four-ridged spines. (G) *Clava multicornis* in situ growing on algae. (H-J) *Bonneviactinia allmanii* gen. nov., comb. nov. (H) growing on *Colus* sp. in situ, imaged by ROV (image credit: MAREANO / Institute of Marine Research). (I) Close-up of fixed medusoid, with visible radial canals and opening; (J) Dense colony on a mollusc shell. (K) *Schuchertinia sarsii* in situ growing on *Pagurus pubescens* (image credit: Bernard Picton). (L-N) *Bonneviactinia arctica* gen. nov., comb. nov.. (L) Small colony growing on a *Mohnia mohni* snail shell. (M) Colony growing on a polychaete tube. (N) Close-up of the complete polychaete tube. (O-P) *Schuchertinia monocarpa*. (O) Two stolonal polyps growing on a leptothecate hydroid. (P) Another colony growing on a different hydroid colony, showing the characteristic large, tube-like spines. Scale bars are approximately 1 mm unless otherwise indicated..

**Table 2.** Morphological and reproductive characteristics of Hydractiniidae species examined in this study. The data include colony morphology, gonophores type, gastrozooid size and tentacle number, spine morphology, distribution, and medusa size, tentacle number, and observed collection season for species with free medusa. Data obtained in this study are indicated by an asterisk (*). Measurements are presented as mean ± SD when multiple specimens were examined; n indicates the number of specimens measured. Literature data are indicated by superscript numbers: (1) Schuchert, P. (2008). The European athecate hydroids and their medusae (Hydrozoa, Cnidaria): Filifera Part 3. Revue Suisse de Zoologie, 115(2), 221–302; (2) Schuchert, P. (2001). Hydroids of Greenland and Iceland (Cnidaria, Hydrozoa) Meddelelser om Grønland, Bioscience 53. Copenhagen, the Danish Polar Center, 2001; (3) Edwards, C. (1972). The hydroids and the medusae *P*. *areolata*, *P. borealis* and *P. carnea*. Journal of the Marine Biological Association of the United Kingdom, 52, 97-144. ND, no data.

| Polyp stage | Type locality | Hydrorhiza | Gonophores | Gastrozooid size [mm] | Number of tentacles | Spines | Distribution |
|---|---|---|---|---|---|---|---|
| ***Bonneviactinia arctica***<br>**(Jäderholm, 1902)** | East Greenland | stolonal | Fixed medusoid, rudimentary tentacles, ring canal, possibly radial canals, up to 10 eggs<sup>(1)</sup> | 0.9 (n=2)* | 5-9 (n=2)* | absent* <sup>(1)</sup> | North Atlantic, Arctic |
| ***Bonneviactinia allmanii***<br>**Bonnevie, 1898** | Faroe Islands | encrusting | Fixed medusoid, rudimentary tentacles, ring canal, radial canals, up to 100 eggs<sup>(1)</sup> | 7 ± 0.4 (n=2)* | 14.8 ± 2.5 (n=2)* | small, 0.1 mm* | Norwegian Sea, Arctic |
| ***Schuchertinia sarsii***<br>**(Steenstrup, 1850)** | Norway | encrusting | Fixed sporosacs, up to 10 eggs<sup>(1)</sup>,<br>no radial canals | 1.6 ± 0.7 (n=5)* | 17 ± 3.5 (n=5)* | smooth, 0.7 – 1.2 mm* | North Atlantic, Norway |
| ***Schuchertinia monocarpa***<br>**Allman, 1876** | Spitsbergen (Svalbard) | encrusting | Fixed sporosacs, up to 50 eggs<sup>(1)</sup>,<br>no radial canals | Up to 3 mm<sup>(2)</sup> | 4 ± 6 (n=2)* | tall, pointed, longer than polyps | Arctic |
| ***Clava multicornis***<br>**(Forsskål, 1775)** | Öresund (Denmark or Sweden) | stolonal/<br>encrusting | Fixed sporosacs, up to 3 eggs<sup>(1)</sup>,<br>no radial canals | 4.3 ± 2.1 (n=7)* | 14.8 ± 4.2(n=7)* | absent | North-Eastern Atlantic<sup>(1)</sup> |
| ***Clavactinia serrata***<br>**Kramp, 1943** | East Greenland | encrusting | Fixed sporosacs, 1 egg<sup>(1)</sup>,<br>no radial canals, spadix present | 1.3 (n=1)*,<br>up to 3<sup>(1)</sup> | 5-9 (n=1)* | short, 0.3 mm*,<br>0.45 - 1.2 mm<sup>(1)</sup> | Arctic<sup>(1)</sup> |
| ***Hydractinia echinata***<br>**(Fleming, 1828)** | British Isles | encrusting | Fixed sporosacs, up to 8 eggs,<br>no radial canals, gonochoric<sup>(1)</sup> | 1.6 ± 0.6 (n=6)*, 5-12<sup>(1)</sup> | 10.8 ± 2.2 (n=6)*, 15-20<sup>(1)</sup> | serrated, 0.5-1 mm (n=6)*,<br>up to 2 mm<sup>(1)</sup> | North-Eastern Atlantic<sup>(1)</sup> |
| ***Podocoryna areolata***<br>**(Alder, 1962)** | UK,<br>Northumberland, | stolonal/<br>encrusting | Medusa (see below) | 0.4 ± 0.1 (n=4)*, up to 2<sup>(1)</sup> | 7.6 ± 2 (n=4)*, 4-14<sup>(1)</sup> | short, smooth, 0.2-0.4 mm (n=4)*,<br>up to 0.9 mm<sup>(1)</sup> | North-Atlantic<sup>(1)</sup> |
| ***Podocoryna borealis***<br>**(Mayer, 1900a)** | USA,<br>Maine, Eastport | encrusting | Medusa (see below) | 1.9 (n=1)*,<br>4-5<sup>(1)</sup> | 8-13 (n=1)*,<br>10-16<sup>(1)</sup> | short, smooth, 0.2 mm, (n=1)*, 0.2-0.6 mm<sup>(1)</sup> | North Atlantic, Maine, Iceland, UK, North Sea, Norway<sup>(1)</sup> |
| ***Podocoryna carnea***<br>**(M. Sars, 1846)** | Florø and Manger, Norway | stolonal/<br>encrusting | Medusa (see below) | 0.6 ± 0.3 (n=11)*, up to 5.5<sup>(1,3)</sup> | 8.9 ± 2.6 (n=11)*,<br>12-19<sup>(1)</sup> | short, smooth, not always present, 0.1-0.2 mm*; 0.2-0.3 mm<sup>(1)</sup> | North Atlantic, UK, Denmark, Sweden, Norway<sup>(1)</sup> |
| **Medusa stage** | **Medusa size** |  | **Number of tentacles** | **Collection seasons** |  |  |  |
| ***Podocoryna areolata*** | 2.8 ± 0.4 (n=6)* |  | >40 | Polyps: April to September; Medusa: March to September |  |  |  |
| ***Podocoryna borealis*** | 2.6 ± 0.5 (n=7)* |  | 16-30 | Polyps: April to October*, November<sup>(1)</sup>; Medusa: April to September |  |  |  |
| ***Podocoryna carnea*** | 1.8 ± 0.1 (n=5)* |  | 4-8 | Polyps: March to September; Medusa: September |  |  |  |

For *Podocoryna,* both polyp and medusa stages were examined, with medusae documented for *P. areolata, P. borealis,* and *P. carnea* (Table 2; Fig. 6). One medusa of *P. carnea* was collected in Norway that resembled *P. exigua*, having only 4 tentacles as described for *P. exigua* (Schuchert, 2008). Freshly collected *Podocoryna* polyps from the type locality were then sampled and reared in laboratory conditions as described in Soto-Angel et al. (2024). This allowed us to monitor development from recently released medusa until day 10, when they showed full maturity (Fig. 7). The medusae were attributed to both *P. carnea* and *P. areolata* based on morphological grounds and later confirmed molecularly. Recently released medusa of *P. carnea* all had 5 or more tentacles (Fig. 7). However, own observations from wild specimens confirm that young *P. carnea* can occasionally have less than 5 tentacles. According to Schuchert et al. 2012, the southern coast of the English Channel is the northernmost distribution for *P. exigua*. No genetically confirmed specimens of *P. exigua* exist outside the Mediterranean, and our data support a distributional difference between these two species.

**Figure 6.**
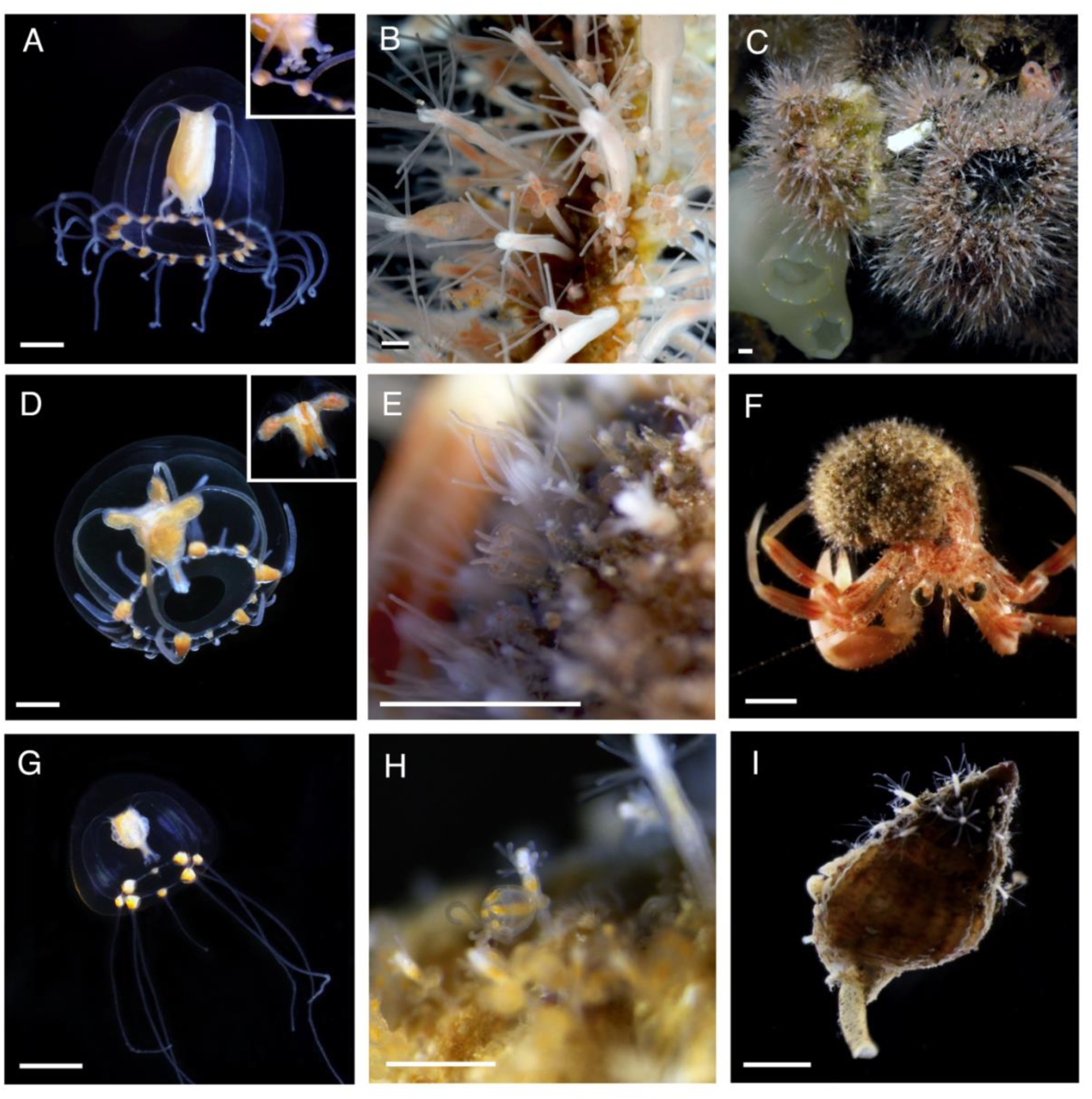
Morphology of medusa-bearing hydractiniid species (*Podocoryna* spp.) observed in this study. (A-C) *Podocoryna borealis*. (A) Mature medusa; inset showing details of the mouth with four, branched oral lips (B) Close-up of a polyp colony with developing medusae. (C) In-situ colony growing on barnacles. (D-F) *Podocoryna areolata*. (D) Mature medusa; inset showing the mouth region with unbranched lips. (E) Close-up of a gastrozooid and a developing medusa, ready to detach. (F) Hermit crab with *P. areolata* colony growing on its shell. (G-I) *Podocoryna carnea*. (G) Mature medusa. (H) Close-up of a gonozooid with medusa attached. (I) *Tritia* sp. mollusk with a small *P. carnea* colony growing on the shell. Scale bars for the medusae (A, D and G) are 1 mm; Scale bars for the other images are approximately 2 mm.

**Figure 7.**
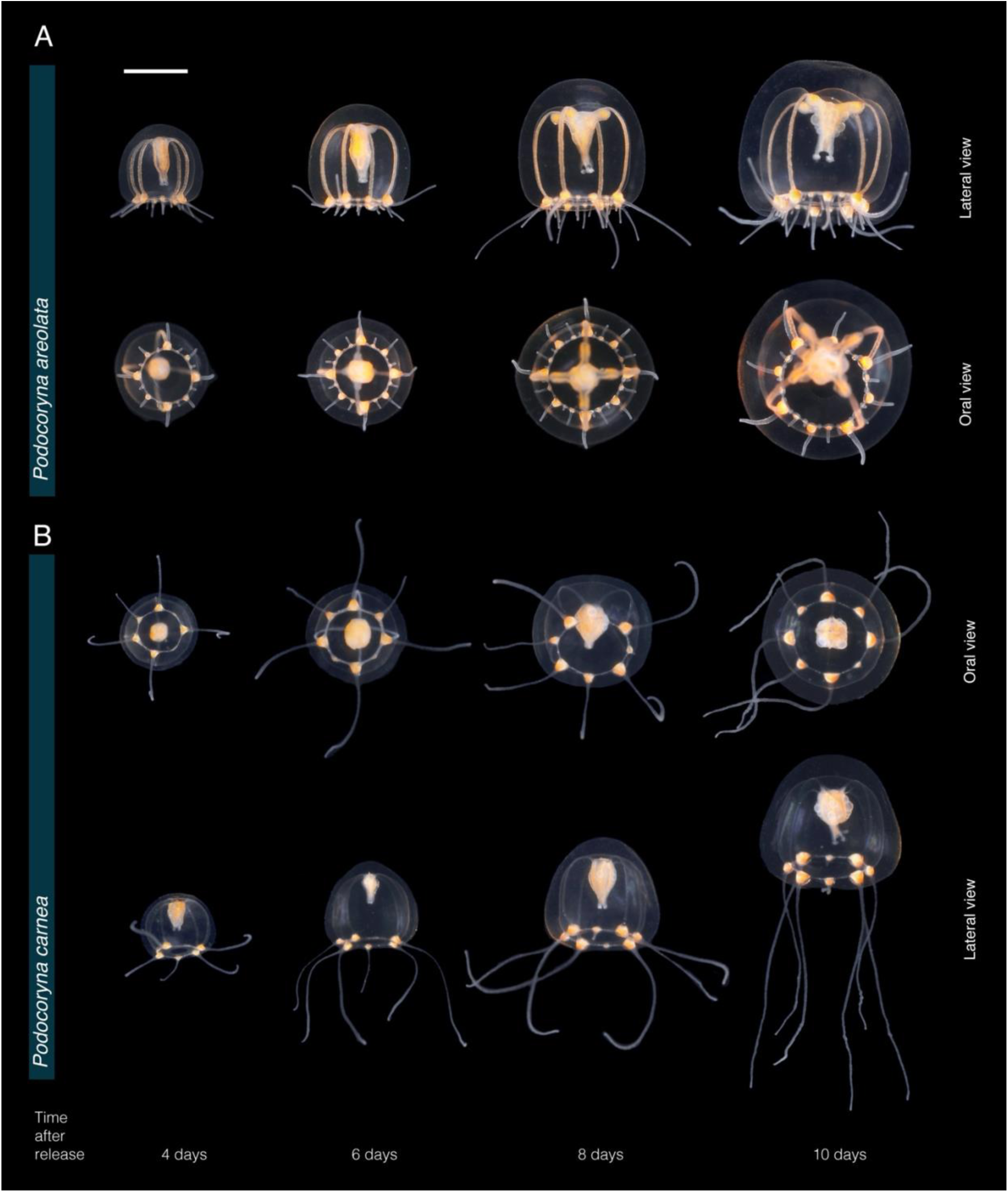
Ontogenetic changes in *Podocoryna areolata* (A) and *Podocoryna carnea* (B) medusae reared under laboratory conditions. Note the difference in size between *P. areolata* (A) and *P. carnea* (B), the differences in the number of tentacular bulbs and tentacles, and the progressive development of gonads, fully mature by day 10 for both species. Scale bar: 1 mm.

### 3.4. Taxonomic account

#### Family Hydractiniidae Agassiz, 1862

##### Bonneviactinia gen. nov

###### Type species

*Hydractinia allmanii* Bonnevie, 1898; here transferred to *Bonneviactinia* gen. nov. as *Bonneviactinia allmanii* (Bonnevie, 1898), gen. nov, comb. nov.

###### Specimens studied

ZMBN 190045; ZMBN 190046; ZMBN 190253; ZMBN 190284 ZMBN 192132; ZMBN 192149; ZMBN 192203, ZMBN 192224 (see Supplementary Table 1 for details); and type material of *Hydractinia allmanii* Bonnevie, 1898 (ZMO_B599 and ZMO_B600).

###### Diagnosis

Hydractiniid hydroids with fixed, non-feeding medusoids retaining a reduced medusa morphology, including a ring canal, radial canals, and rudimentary marginal tentacles. Gastrozooids and colony morphology variable. Gonozooids reduced, typically bearing two gonophores, as medusoids. *Bonneviactinia* is distinguished from morphologically similar hydractiniid genera by its molecularly supported monophyletic lineage recovered in mitochondrial 16S, concatenated 16S + 18S + 28S, and mitochondrial-genome phylogenies.

###### Included species

*Bonneviactinia allmanii* (Bonnevie, 1898), gen. nov., comb. nov. and *Bonneviactinia arctica* (Jäderholm, 1902), gen. nov., comb. nov.

###### Etymology

*Bonneviactinia* is named in honour of the Norwegian zoologist Kristine Bonnevie (1872-1948), who made important contributions to the study of marine invertebrates and described *Hydractinia allmanii* Bonnevie, 1898, the type species of this genus. The suffix -actinia refers to its placement within Hydractiniidae and follows the naming tradition of several genera within the family.

###### Remarks

The genus is supported as a distinct lineage in mitochondrial 16S, concatenated 16S + 18S + 28S, and mitochondrial-genome phylogenies (Figs. 2-4). The two included species form a strongly supported lineage across all analyses. *Bonneviactinia* gen. nov. is morphologically heterogeneous in colony form. *Bonneviactinia allmanii* gen. nov., comb. nov., forms dense colonies composed by large polyps with a coalesced stolonal plate and usually bears small hydrorhizal spines, whereas *B. arctica* has a loose, non-coalesced stolonal hydrorhiza without spines. They nevertheless share fixed medusoids retaining radial canals and rudimentary marginal tentacles. The presence of radial canals is not unique to *Bonneviactinia* gen. nov. and also occur in other hydractiniids with medusoid gonophores, including species assigned to *Stylactis* (Schuchert 2008). The distinctness of *Bonneviactinia* is therefore primarily supported by the molecular data, in which *B. allmanii* and *B. arctica* form a strongly supported lineage across mitochondrial and nuclear datasets. Notably, *B. arctica* was originally described as *Stylactis arctica* Jäderholm, 1902, reflecting its loose stolonal colony, absence of hydrorhizal spines, and reduced gonophores retaining radial-canals. These features also resemble those of species assigned to *Stylactis*. In our molecular analyses, however, *B. arctica* does not form a clade with *Stylactis*, supporting its placement in a separate genus despite these morphological similarities. This provides a new example of how similarities in colony and reproductive morphology do not necessarily reflect generic relationships in Hydractiniidae.

##### *Bonneviactinia allmanii* (Bonnevie, 1898), gen. nov., comb. nov

###### Original combination

*Hydractinia allmanii* Bonnevie, 1898. **Synonymy:** *Hydractinia ornata* Bonnevie, 1898

###### Material examined

Type material (ZMO_B599 and ZMO_B600); ZMBN 190045; ZMBN 190046; ZMBN 190253; ZMBN 192149; ZMBN 192224 (full collection details in Supplementary Table 1)

###### Remarks

Examination of the type material (ZMO_B599 and ZMO_B600) provides a morphological reference for the identification of the specimens assigned to *B. allmanii* in this study. These specimens provide the first molecular data for the species and place it in a strongly supported lineage together with *B. arctica*.

##### *Bonneviactinia arctica* (Jäderholm, 1902), gen. nov., comb. nov

###### Original combination

*Stylactis arctica* Jäderholm, 1902. **Synonymy:** *Stylactaria arctica* (Jäderholm, 1902)

###### Material examined

ZMBN 190284; ZMBN 192132; ZMBN 192203 (full details in Supplementary Table 1)

###### Remarks

Originally described as *Stylactis arctica* Jäderholm, 1902, this species was subsequently assigned to *Hydractinia*. Its original placement in *Stylactis* is morphologically understandable given its loose stolonal hydrorhiza, lack of hydrorhizal spines, and reduced medusoids. However, our molecular analyses place *B. arctica* together with *B. allmanii* rather than with the *Stylactis* species of previous studies, supporting its transfer to *Bonneviactinia*.

##### *Schuchertinia sarsii* (Steenstrup, 1850), comb. nov

###### Original combination

*Podocoryne sarsii* Steenstrup, 1850. **Synonymy:** *Podocoryna sarsii* Steenstrup, 1850

###### Material examined

ZMO_B686 and ZMO_1347 (type specimens); ZMBN 185345, ZMBN 185316, ZMBN 199643, ZMBN 199644, ZMBN 185306, ZMBN 185308, ZMBN 185309, ZMBN 185521 (full details in Supplementary Table 1)

###### Remarks

Specimens identified as *S. sarsii* in this study group with sequences previously published under the name *Schuchertinia allmanii* (Miglietta et al., 2009; Miglietta & Cunningham, 2012). Comparison with the type material, together with the molecular results, supports their identification as *S. sarsii* rather than *B. allmanii*. These findings indicate that the previously published specimens were likely misidentified and establish a molecular reference for *S. sarsii*.

##### *Schuchertinia monocarpa* (Allman, 1876), comb. nov

###### Original combination

*Hydractinia monocarpa* Allman, 1876

###### Material examined

ZMBN 199611, ZMBN 199627 (full details in Supplementary Table 1)

###### Remarks

The specimens identified as *Hydractinia monocarpa* in this study exhibit the long, pointed hydrorhizal spines and gonozooids devoid of tentacles and 1-2 sporosacs characteristic of the species (Schuchert, 2008). Their placement within *Schuchertinia* clade II provides the first molecular evidence supporting the assignment of the species to this genus. However, because the monophyly of *Schuchertinia* remains unresolved, this generic placement should be regarded as provisional pending broader molecular sampling, particularly of the type species and other incompletely characterized members of the genus.

### 3.5 Host and substrate associations

Host associations varied among the hydractiniid species examined, with several species showing distinct patterns of association with particular host types (Fig. 8). Molecular confirmation of species identities allowed these associations to be assessed at the species level.

**Figure 8.**
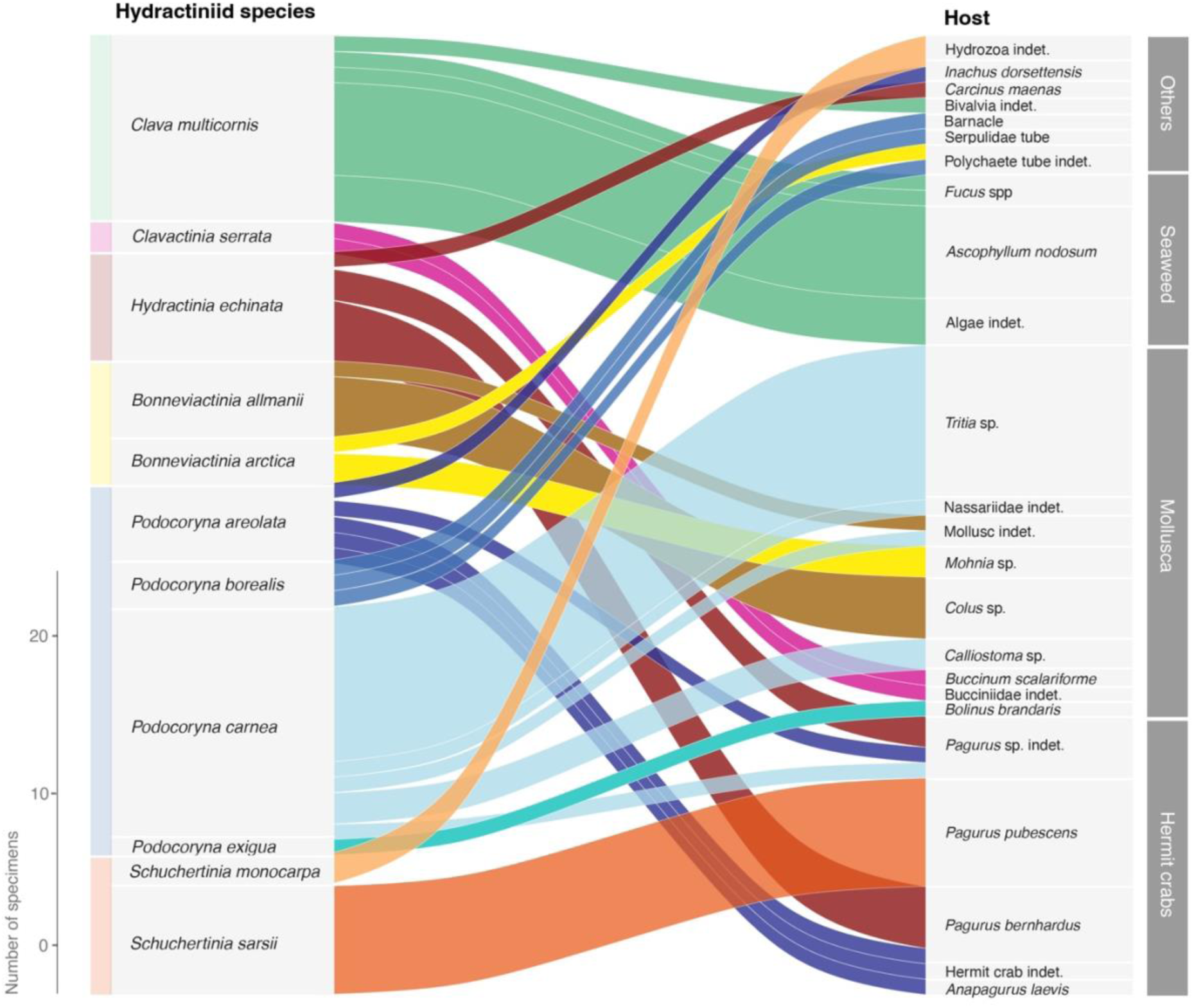
Host and substrate associations of Hydractiniid polyps. Alluvial plot showing associations between Hydractiniid species and recorded host types. Hosts are grouped into broad categories according to host identity, and flow widths represent the number of specimens associated with each host. Flows are coloured by Hydractiniid species.

*Clavactinia serrata* was predominantly associated with *Buccinum* spp. (Buccinidae). Colonies were found on shells inhabited both by the gastropod and by hermit crabs, suggesting that the association was primarily with the shell rather than with the organism inhabiting the shell. *Hydractinia echinata* was found exclusively on gastropod shells inhabited by *Pagurus* hermit crabs, predominantly *Pagurus bernhardus*, and was not recorded on other host types.

*Bonneviactinia allmanii* gen.nov., comb.nov. was exclusively found on mollusc shells inhabited by their molluscan hosts, including *Colus* sp. In contrast, *Bonneviactinia arctica* gen.nov, comb.nov. was recorded from two distinct substrate/host types. One colony was growing on a muddy polychaete tube; examination of a large collection of polychaete tubes from the region did not reveal additional colonies. Other colonies were found on the mollusc *Mohnia* sp.. These observations indicate that *Bonneviactinia arctica* gen.nov., comb.nov may have relatively broad host or substrate associations.

*Podocoryna areolata* polyp colonies were recorded from a variety of hosts, including the spider crab *Inachus dorsettensis* and several hermit crab species, including *Pagurus* spp.*, P. bernhardus,* and *Anapagurus laevis.* The species was predominantly associated with crabs rather than snails. *Podocoryna borealis* polyp colonies were found on several benthic organisms that were fixed or sessile and provided a relatively large surface for attachment, including fields of barnacle shells, Serpulidae, and other polychaete tubes. The observed associations therefore largely comprised immobile substrates.

*Podocoryna carnea* was also recorded from several host types but was most frequently found on *Tritia* sp. inhabited by the snail itself. Additional colonies were found on other gastropods, including *Calliostoma* sp. and other Nassariidae, while one colony was found on *Pagurus* sp., where the colony was associated with a shell inhabited by a hermit crab. Thus, *P. carnea* was predominantly associated with gastropod shells, although host association alone cannot be used to distinguish *P. carnea* from *P. areolata. Schuchertinia monocarpa* was exclusively found growing on other hydrozoans (Leptothecata), encompassing different host species (e.g. *Halecium* sp). *Schuchertinia sarsii* was also recorded exclusively from one host association, on shells inhabited by *Pagurus pubescens*.

## 4 Discussion

Despite being one of the best-studied hydrozoan families, the diversity and taxonomy of Hydractiniidae in the North Atlantic remain incompletely resolved (Schuchert, 2001; 2008; 2012). This is unexpected given the prominence of *Hydractinia echinata* and *Podocoryna americana* (previously identified as *P. carnea)* as model organisms in developmental biology, regeneration and life-cycle evolution (Weismann, 1883; Plickert et al., 2012; Sanders et al., 2018; Galliot & Schmid, 2002). At the same time, the family has long presented difficulties for species identification because of the paucity of diagnostic morphological characters, morphological plasticity, and the occurrence of cryptic lineages (Cartwright & Nawrocki, 2010; Schuchert, 2012; Miglietta et al., 2009). The characteristic, fur-like colonies growing on gastropod shells have been familiar to beachcombers and represented in coastal identification guides, while several less conspicuous species have received comparatively little attention. An integrative approach combining morphological and molecular data is particularly valuable in this group, both for resolving species boundaries and for associating morphologically different life stages, such as hydroid colonies and medusae. Here, we resolved 10 hydractiniid species spanning shallow coastal habitats to deep-sea environments below 2500 m. Our results clarify the identities and generic placement of several historically problematic taxa, including *Schuchertinia sarsii* comb. nov., *Bonneviactinia allmanii* gen. nov., comb. nov., and *Bonneviactinia arctica* gen. nov., comb. nov. We show that *S. sarsii* and *B. allmanii* belong to separate genera, while *B. allmanii* and *B. arctica* form part of the same evolutionary lineage. In regard to their life cycles, both species develop reduced medusoids, with rudimentary tentacles and canal systems. In addition to resolving long-standing taxonomic issues, we provide vouchered DNA barcodes for all examined species and the first complete mitogenomes for seven species Together, these data provide a reference framework for future species identification and diversity assessments, including both traditional morphological surveys and molecular approaches such as DNA barcoding and metabarcoding/eDNA. In particular, linking sequence data to morphologically identified, vouchered specimens provides an essential reference for interpreting sequence-based surveys, while documented geographic distributions help predict which taxa may occur in particular habitats or regions.

### 4.1 Diversity and distribution of Hydractiniidae in the NE Atlantic

Several hydractiniid species currently regarded as broadly distributed may have more restricted distributions than historical records suggest. For example, *Hydractinia echinata* was recorded as far north as the Arctic Ocean (Schuchert, 2008), but our findings place its northern limit in southern Norway. *H. echinata* is well documented from northwestern Europe and southwestern Scandinavia, and previous records from further north therefore warrant particular scrutiny. One possible explanation for apparently broader historical distributions is confusion with morphologically similar hydractiniids. In northern Norway and towards Svalbard, some historical records of *H. echinata* may therefore have been misidentified, particularly as *S. sarsii* and *Clavactinia serrata* occur in the region. The apparent rarity of *S. sarsii* in the historical record may partly reflect such taxonomic confusion.

The original description of *S. sarsii* is based on a sample collected in southwestern Norway by Michael Sars (Sars, 1846), who identified the colony as *Podocoryna carnea.* A few years later, Johannes Steenstrup (in Lütken, 1850) reexamined this specimen, concluding that it represented the new species *Hydractinia sarsii.* The species has only been recorded a few times after its original description from the Oslofjord, the Kattegat close to Gothenburg but also Svalbard, Greenland and Iceland (Jägerskiöld, 1971; Christiansen, 1972), and Schuchert (2008) noted that its biology remains poorly known. Our molecular data demonstrate that *S. sarsii* is much more widespread than the published occurrence records alone would suggest, including regions where *H. echinata* was not detected in our study (i.e. Svalbard). This highlights how incomplete taxonomic knowledge can translate directly into apparently anomalous species distributions, and our molecular and morphological data provide an opportunity to disentangle this historical confusion and establish a reference identity for these species.

The only lineage in our dataset predominantly associated with offshore and deeper-water habitats is the new genus *Bonneviactinia*. *Bonneviactinia allmanii* gen.nov.,comb.nov. occurred in the Norwegian Sea and around Svalbard at approximately 250 to >1000 m depth, with records following the continental shelf margin towards its type locality in the Faroe Islands. *Bonneviactinia arctica* gen.nov., comb.nov was similarly restricted to offshore habitats but occurred at substantially greater depths, from approximately 700 to >2500 m, around Svalbard. Our sampling documented three additional species occurring in the Arctic: *Clavactinia serrata, S. sarsii,* and *S. monocarpa. Clavactinia serrata* was found only in Svalbard, where it occurred relatively frequently, including at depths shallower than 100 m. This agrees with its previously attributed Arctic distribution (Schuchert, 2008; Ronowicz et al. 2015). *S. monocarpa* was likewise restricted to Svalbard in our dataset, corresponding to its type locality, whereas *S. sarsii* showed a substantially broader boreoarctic distribution.

The three species of the medusa-bearing genus *Podocoryna* reported here occurr sympatrically in southern Norway. *Podocoryna carnea* appeared to have the most restricted distribution, with no specimens detected north of Trondheim, whereas medusae of *P. areolata* and *P. borealis* were also recorded from the northernmost waters of mainland Norway. *P. borealis* was found in depths down to 1250 m in the fjords. Neither hydroid colonies nor medusae of *Podocoryna* were detected offshore or around Svalbard. Our records extend the documented distributions of *P. areolata* and *P. borealis* into Arctic waters, from which they have not previously been reported in the Norwegian region (Schuchert, 2009; Edwards, 1972). Historical literature already indicates that the distribution of *P. carnea i*s concentrated in northwestern Europe. Morphological identification of the hydroid-stage of *Podocoryna* species can be problematic due to the lack of diagnostic characters. This is especially true for *P. carnea* and *P. areolata:* we noted that medusa-bearing hydroids growing on a shell inhabited by a mollusc (such as *Tritia* sp) are usually *P. carnea*. Conversely, colonies growing on hermit crabs (*Pagurus* sp.) cannot easily be identified, as both species can grow on hermit crabs (Fig. 8). Here, molecular identification is necessary to identify reliably to species.

Two species are reported to occur in the study area but were not found in the present study: *Hydractinia ingolfi* and *Hydractinia carica. Hydractinia ingolfi* is known from very few records and was originally described from deep-sea material growing on the ophiurid *Homalophiura tesselata* at approximately 2100-3200 m depth. Schuchert (2008) considered *H. ingolfi* unlikely to be conspecific with *H. arctica*, emphasizing its unusual association with a deep-sea ophiurid and its characteristic large, trumpet-shaped hypostome. In our study, no deep-sea brittle stars bearing hydractiniid colonies were found, despite intense search in the existing museum material from the region. The species could simply be very rare, but the similar depth range and morphology of our *Bonneviactinia arctica* gen.nov., comb.nov. material, together with the diverse substrates colonized by *B. arctica* gen.nov., comb.nov., raises the possibility that some historical deep-sea records may have been mistakenly assigned to *H. ingolfi.* However, this cannot be resolved without molecular evidence for the species. The type locality of *H. ingolfi* lies far from our sampling area, at 58°20′N, 40°48′W south of Greenland. Thus, its absence from our material does not rule out the possibility that it is a distinct, poorly recorded species. The present data therefore support retaining *H. ingolfi* as an unresolved taxon rather than treating it as conspecific with *Bonneviactinia arctica* gen.nov., comb.nov.

The lack of a sequenced representative of *Hydractinia carica* is difficult to interpret because the species has a complicated taxonomic history. It has previously been treated under the name *H. minuta*, published descriptions are inconsistent, and it has been considered morphologically similar to *H. monocarpa*, all of which complicate the evaluation of historical records. In our material, specimens identified as *H. monocarpa* had the long, pointed spines characteristic of that species, whereas the shorter, smooth spines described for *H. carica* by Schuchert (2008) were not observed. We obtained one putative *H. carica* specimen (NOAH0373), but sequencing was unsuccesful, and we recovered no molecular lineage corresponding to a convincing morphological match. Historical records of *H. carica* may nevertheless include other morphologically similar hydractiniids. In particular, *H. carica* and *Clavactinia serrata* overlap geographically and have both been reported from bucciniid shells; because *C. serrata* was also identified in our material from the northern part of the study area, misidentification is one possible explanation for some historical records of *H. carica.* This remains to be tested, as we have no evidence for them to be conspecific. Resolving its status will require targeted sampling in Svalbard and the northern Norwegian and Arctic seas, including reproductive colonies and molecular data linked to well-documented vouchers.

### 4.2 Life-cycle evolution and biosystematics

Hydractiniids span nearly the full range of hydrozoan life cycles, from species releasing free-swimming feeding medusae, through liberated or fixed non-feeding eumedusoids retaining radial canals, to fixed sporosacs with few vestiges of medusa morphology (Miglietta & Cunningham, 2012). Medusa reduction has been suggested to be the consequence of paedomorphic progenesis, in which sexual maturation is accelerated relative to medusa development, giving rise to reduced, medusa-like gonophores capable of sexual reproduction prior to liberation (Gould 1977; Boero and Sara 1987; Cunningham & Buss, 1993; Miglietta & Cunningham, 2012). Loss of the feeding medusa has been inferred repeatedly in Hydractiniidae, and the degree of medusa reduction has been interpreted as a developmental modification rather than a phylogenetically informative character (Miglietta & Cunningham, 2012). In particular, the nested position of medusa-bearing *Podocoryna* within lineages lacking feeding medusae illustrates the evolutionary complexity of this character and has previously led to competing hypotheses concerning repeated loss versus regain of the feeding medusa (Miglietta & Cunningham, 2012). Ancestral state reconstructions by Miglietta and Cunningham (2012) implied the re-emergence of the free medusa stage after more than 100 million years of absence, although the authors favoured the scenario of multiple independent losses.

Here, we provide new evidence for reassessing the evolution of life-cycle strategies in a family in which developmental characters have historically been closely intertwined with taxonomy. When life cycles strategies are mapped onto the newly produced phylogeny, they reveal a pattern that can be interpreted in two possible ways: a stepwise gain of medusa characters in Hydractiniidae, or as repeated, independent losses of the medusa stage across several hydractiniid lineages, as favoured by prior authors (Miglietta & Cunningham, 2012). Fixed sporosacs represent the most reduced condition and occur in several of the more distantly related lineages, whereas *Stylactis* and *Bonneviactinia* retain radial canals in (presumably) fixed medusoids (Fig. 9). Further putative retention of medusa morphology is seen in *Bouillonactinia*, which produces (mostly) liberated medusoids, while *Podocoryna* retains (or regains) a fully developed, free-swimming feeding medusa. Although this pattern should not be interpreted as a single evolutionary sequence without formal ancestral-state reconstruction, it potentially illustrates the repeated modification of the medusa stage across the family, with successive retention of increasingly complex medusa characters in some lineages.

**Figure 9.**
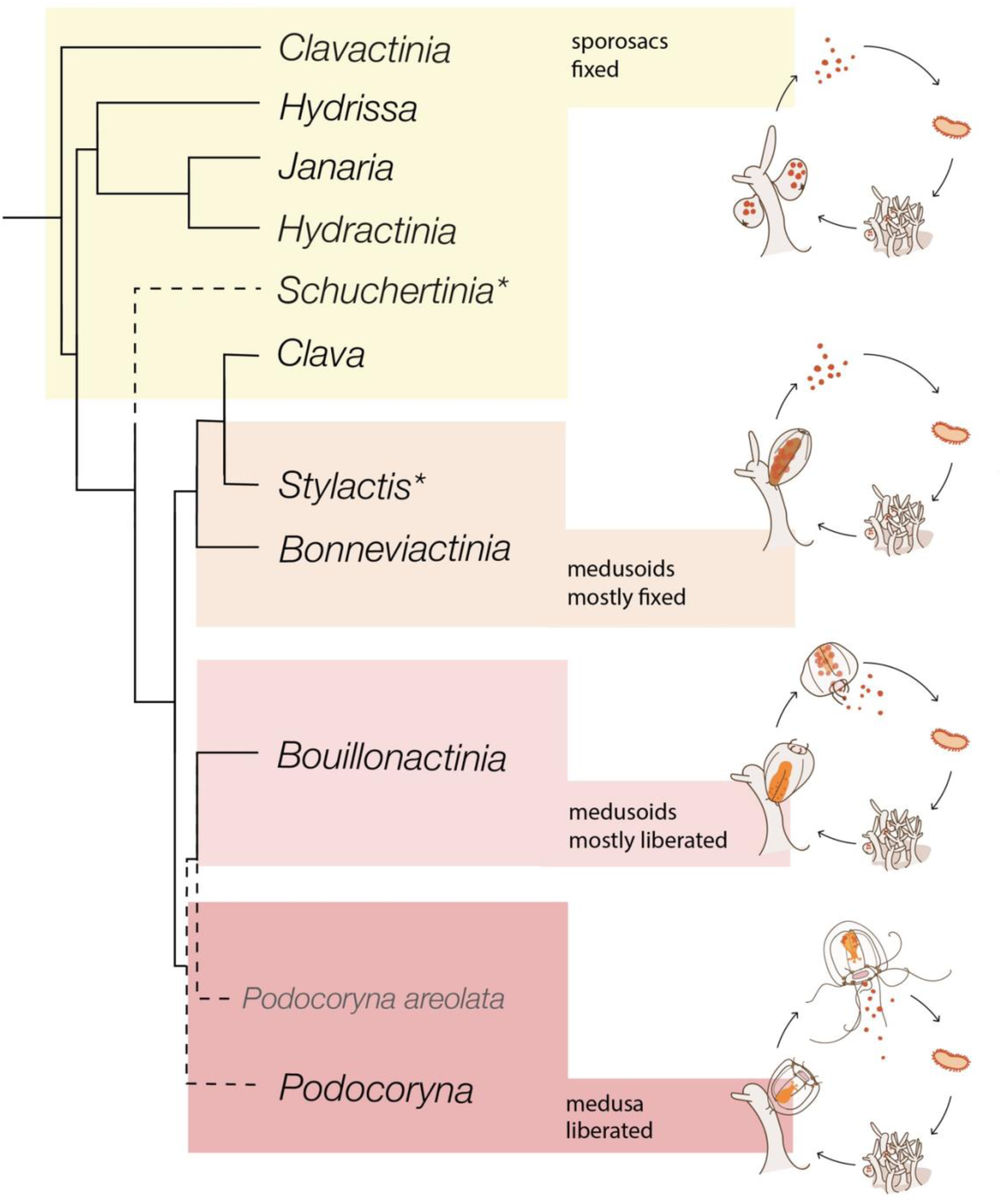
Simplified phylogeny of the family Hydractiniidae with selected genera collapsed (based on our concatenated and 16S phylogeny. Any unresolved relationships are indicated with dashed line. Reproductive strategies are indicated for each genus where known. The asterisks indicates conflicting information about reproductive structures and/or different gonophores as indicated. For *Schuchertinia* two species in the genus have different gonophores reported: *Schuchertinia antonii* is described to develop radial canals and gonophores called cryptomedusoids in its original description (Miglietta, 2010); and *S. epichoncha*, has reported eumedusoids bearing radial canals and rudimentary tentacles (Stechow, 1908; Hirohito, 1988). Also note that the genus *Schuchertinia* was paraphyletic, and the placement of the genus is not resolved for now. Especially *S. epichoncha* was not resolved within the genus (See Figs. 2 and 3, and Supplementary Figure 2). For *Stylactis* spp. reproductive information taken from Schuchert (2008), with radial canals in *S. inermis* but none in *S. fucicola*, but both are described as fixed.

Our results refine previous findings in several ways. On one hand, the specimens misidentified as *Schuchertinia allmanii* in previous works (Miglietta et al., 2009; Miglietta & Cunningham, 2012) correspond to *S. sarsii* comb. nov., which bears fixed sporosacs, clarifying the *Bonneviactinia allmanii* gen. nov, comb. nov., sequenced here for the first time, forms a distinct lineage together with *B. arctica*, characterised by fixed medusoids (Tab. 1). The species of *Schuchertinia* for which the whole mitogenome and complete 18S and 28S nuclear markers are available are characterized by the presence of fixed sporosacs. In this regard, our results show that life-cycle strategies may be more uniform within genera than previously thought. However, the type species of *Schuchertinia*, *S. epiconcha*, releases eumedusoids. The weak support for the monophyly of *Schuchertinia,* together with the long branch separating *S. epiconcha* from its sporosac-bearing congeners, may indicate the presence of an additional, undescribed genus. The lack of complete nuclear markers and mitogenomes for most species of *Schuchertinia*, including its type species, precludes a formal reassessment of the genus with the available evidence.

On the other hand, *Bonneviactinia* falls within a strongly supported clade that also includes *Clava, Podocoryna* and *Bouillonactinia.* This clade spans the full range of gonophore types found in the family: fixed sporosacs in *Clava*, fixed medusoids in *Bonneviactinia* and presumably *Stylactis,* liberated eumedusoids in *Bouillonactinia*, and feeding medusae in *Podocoryna* (Fig. 9). This pattern is consistent with extensive modification and repeated reduction of the medusa stage throughout Hydractiniidae, rather than with a simple unidirectional trajectory towards the occurrence of a free medusa.

Our results agree with Miglietta & Cunningham (2012) in recovering *Bouillonactinia* and *Podocoryna* as sister groups. However, when complete nuclear markers are included and *Podocoryna areolata* is added to the analysis, *Podocoryna* is recovered as paraphyletic, with *P. areolata* sister to *Bouillonactinia* (83/59) rather than to the remaining species of the genus. Support for this relationship is weak and the taxonomic representation of *Bouillonactinia* is limited to a few species with partial sequences (though even in the chimeric analysis, the support was weak), with no mitogenome available for the genus. The paraphyly of *Podocoryna* should therefore be regarded as provisional. If confirmed, however, it has direct implications for the evolutionary history of the free medusa stage within Hydractiniidae. Either the free feeding medusa was lost in *Bouillonactinia*, or it arose independently in *P. areolata* and in the remaining *Podocoryna*. The first scenario requires a single transition and involves only a partial reduction, since all molecularly confirmed species within *Bouillonactinia* release swimming eumedusoids with radial canals, consistent with truncation of medusa development. The second requires two independent re-gains of a complex structure. The first scenario is therefore considerably more parsimonious, and it is consistent with the repeated reduction of the medusa observed across the family, including within *Podocoryna* itself (e.g. in *P. pruvoti*; Miglietta & Cunningham, 2012). Resolving the relationships among *Podocoryna* species and increasing *Bouillonactinia* taxonomic coverage and molecular resolution will be essential to determine whether *Podocoryna* as currently defined is monophyletic, and therefore the evolutionary lability of the feeding medusa stage within Hydractiniidae.

Life-cycle and colony traits have previously been associated with host type, with loss of the feeding medusa linked to host specialization and hermit-crab specialists consistently bearing sporosacs and forming encrusting colonies (Miglietta & Cunningham, 2012). Our results suggest that this relationship is not universal across Hydractiniidae. *Clavactinia* combines fixed sporosacs with encrusting colonies on molluscan shells, whereas *Bonneviactinia,* despite having a reduced, fixed medusoid stage, occurs on a broader range of substrates, including molluscs and polychaetes. This contrast suggests that gonophore evolution and host specialization are not necessarily coupled and that similar reductions in the medusa may occur under different ecological circumstances. Host use should therefore be considered alongside, rather than as a direct correlate of life-cycle evolution in Hydractiniidae.

### 4.3 From model organisms to reference resources

The new taxonomic framework provided here is particularly relevant for Hydractiniidae because several species in the family are now consolidated model organisms. *Hydractinia echinata* has a long history as model for developmental biology, stem cells, regeneration and allorecognition, while *Podocoryna* has been widely used to investigate colony organization, life-cycle evolution and developmental processes (Miglietta & Cunningham, 2012; Frank et al., 2020). Because model species underpin comparative and experimental work, accurate identification is essential: findings on development, physiology and life history are attributed to a named species and then generalized, so misidentification can propagate errors across studies (Miglietta, 2023). This issue is not unique to Hydractiniidae: cryptic diversity and taxonomic uncertainty have also been demonstrated in other marine model organisms, emphasizing that taxonomic verification is an important component of reproducibility and comparability among studies (Sigwart et al., 2023). Within Hydractiniidae, where closely related species can differ in colony morphology, reproductive structures and life-cycle configuration, molecularly supported identifications can therefore provide an important foundation for interpreting and comparing experimental results.By linking genetically characterized specimens to morphological identifications, this study provides voucher-backed reference material against which future specimens and sequence data can be compared. This is particularly valuable in Hydractiniidae because several species share broadly similar colony morphologies, whereas diagnostic reproductive structures may vary among species and may not be present in all specimens or life-cycle stages. DNA barcodes can therefore complement morphological identification by providing an independent means of confirming species assignments, while complete mitochondrial genomes provide a multi-gene molecular reference for phylogenetic and comparative genomic studies. Importantly, the value of such resources depends not only on the sequence itself but also on the quality and traceability of the associated taxonomic information. Reference-library guidelines emphasize that reliable barcodes should be linked to accurately identified specimens, vouchers and metadata, allowing sequence records to be reassessed as taxonomy changes (Rimet et al., 2021). In this respect, the combination of morphological documentation, locality data, voucher material and sequence data generated here provides a more durable reference than an isolated sequence record without a verifiable taxonomic basis.

These resources are also directly relevant to metabarcoding approaches, including environmental DNA (eDNA). Molecular surveys identify unknown sequences through comparison with reference databases, meaning that incomplete taxonomic coverage can prevent species-level assignments or result in sequences being assigned only to higher taxonomic levels. The importance of this limitation has long been recognized for marine metabarcoding, for which reference sequences from accurately identified species are considered essential for reliable molecular taxonomic assignment (Bucklin et al., 2021). Recent regional efforts further illustrate the value of targeted reference-library development. In the Gulf of Alaska, Questel et al. (2025) generated 1,462 sequences representing 254 taxa, including 107 species for which no publicly available sequence data had previously been available, among them 12 undescribed cnidarian taxa. Similarly, the recent GEANS initiative generated 4,005 COI barcodes from 715 North Sea macrobenthic species, while still representing only around 29% of the estimated regional fauna, demonstrating both the utility of coordinated reference-library development and the substantial remaining taxonomic gap (Christodoulou et al., 2025).

Hydrozoans are particularly relevant in this context because they can be readily detected in molecular surveys of marine communities (Berry et al., 2019). However, detection depends on both the sampling approach and the biology of the target organisms: zooplankton metabarcoding and eDNA recover different components of the hydrozoan community (Ohnesorge et al., 2023), and water-column eDNA does not directly represent benthic assemblages (Antich et al., 2021). Improved reference databases are therefore necessary for interpreting molecular surveys, but cannot by themselves eliminate other sources of variation, such as DNA transport, shedding rates, primer bias and marker resolution.

In this sense, the resources generated here extend beyond the specific findings presented in this study. The Norwegian and broader North Atlantic fauna includes hydractiniids occupying shallow coastal, offshore and deeper-water habitats, and our results suggest that some historical occurrence records should be interpreted with greater taxonomic caution. Expanded molecular reference coverage will allow future barcoding, metabarcoding and eDNA surveys to test such records and to assign otherwise unidentified sequences with greater confidence. This is relevant for inconspicuous or morphologically similar species, which conventional biodiversity surveys usually underepresent. In this context, the value of a reference barcode goes beyond the sequence of a given species; it anchors that sequence to a specimen whose morphology, locality and taxonomic identity are documented. Building such voucher-linked resources for Hydractiniidae will therefore support not only future species identification, but also comparative developmental studies, phylogenetics, biodiversity assessments, and molecular monitoring of marine communities.

## Supplementary Material

- Supp File 1: 16S tree worldwide
- Supp File 2: Figure of 16S+28S+18S tree with chimeras (*Bouillonactinia* gen. nov with 16S from a different specimen, which were missing 16S in the other tree); also includes *Clavactinia serrata* (chimera of our two specimens, complementing the existing sequences); and *Schuchertinia epichoncha* (which only has short 28S, no chimera possible).
- Supp Table 1: all specimens sampled or analyzed for this study (sequenced or morphological analysis) + metadata
- Supp Table 2: additional sequences (mined from NCBI; worldwide are not included in this table yet) + metadata if available

## Acknowledgments

We thank Louise Lindblom and Vincent McDaniel for support with DNA laboratory work and Mariam Rezapoor for assistance with sampling. We are grateful to Nicolas Straube for assistance with library preparation protocols. We thank Marta Gil, Bernard Picton and Christin Morrow for assistance with sampling and imaging. We are grateful to Matthew Travert for information on *Podocoryna* sp. from the USA laboratory strain and for providing sequence data. We want to express our gratitude to Mareano and OceanX for facilitating the collection of *S. allmanii* specimens and for providing ROV imagery. We also thank Piotr Bałazy for providing hydractiniid specimens collected during several diving expeditions to Spitsbergen and Norway, and Maciej Mańko for assistance with mitochondrial genome analyses and interpretation. Special thanks go to Daniel Chorrout, Pawel Burkhardt and Anne Aasjord at the Michael Sars Centre (UiB), which supported the short-term culture of *Podocoryna* spp. at the Appendicularian Facility; as well as Miguel Candelas (Oceanografic València) and Borja Mercado (Universidad Europea de Valencia) for facilitating access to specimens of *Podocoryna exigua.* We gratefully acknowledge the Norwegian Barcode of Life (NorBOL) for supporting the sequencing associated to the Norwegian Taxonomy Initiative projects (see funding section).

## Author contributions

LMB: Conceptualization; Methodology; Formal analysis; Investigation; Software; Visualization; Writing - Original Draft

AH: Validation; Resources; Funding acquisition; Writing – Review & Editing

MR: Resources; Funding acquisition; Writing – Review & Editing

LM: Conceptualization; Methodology; Resources; Supervision; Funding acquisition; Writing – Review & Editing

JJSA: Conceptualization; Methodology; Validation; Resources; Funding acquisition; Writing – Review & Editing

