## Supplementary figures and images for "Revisiting Hydractiniidae (Hydrozoa): Mitogenomic Insights into the Diversity, Systematics and Life-Cycle Evolution of North Atlantic and Arctic Species"

### Supplementary File 1

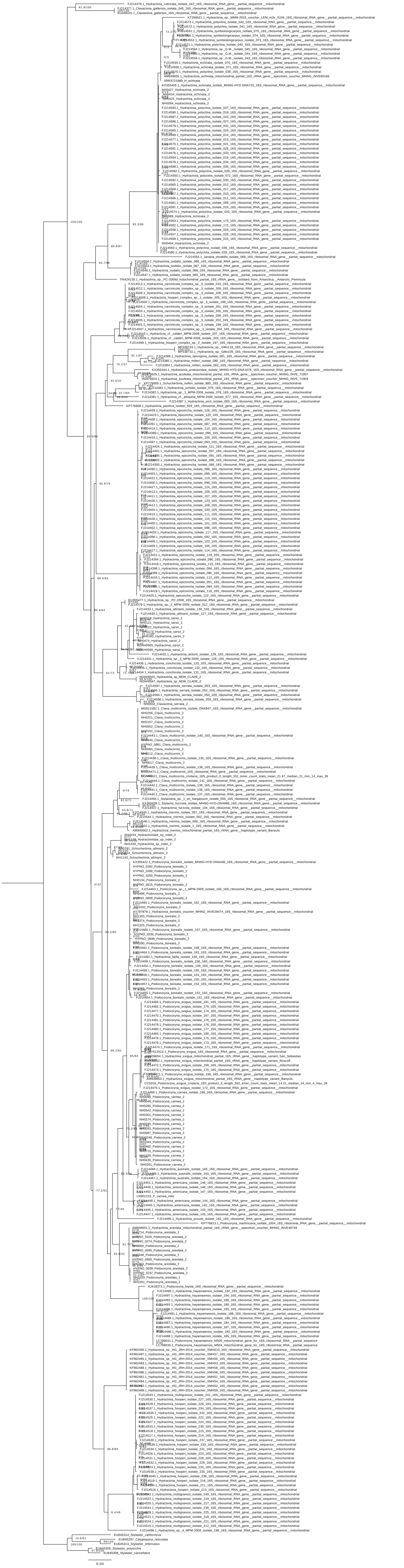
