## Supplementary File 2 for "Revisiting Hydractiniidae (Hydrozoa): Mitogenomic Insights into the Diversity, Systematics and Life-Cycle Evolution of North Atlantic and Arctic Species"

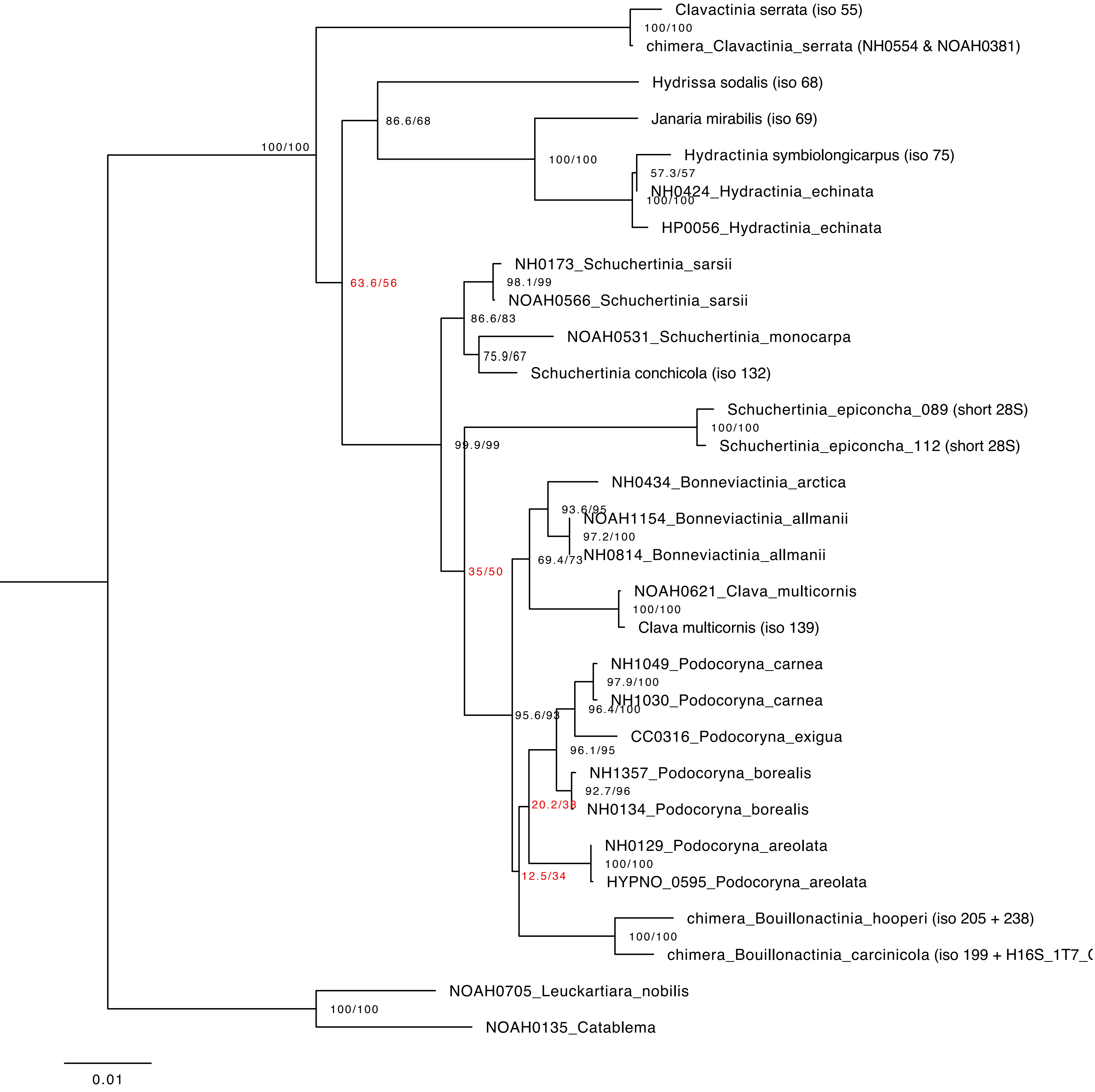

Clavactinia serrata (iso 55)
100/100
chimera_Clavactinia_serrata (NH0554 & NOAH0381)
Hydrissa sodalis (iso 68)
86.6/68
Janaria mirabilis (iso 69)
100/100
Hydractinia symbiolongicarpus (iso 75)
57.3/57
NH0424_Hydractinia_echinata
100/100
HP0056_Hydractinia_echinata
100/100
NH0173_Schuchertinia_sarsii
98.1/99
NOAH0566_Schuchertinia_sarsii
63.6/56
86.6/83
NOAH0531_Schuchertinia_monocarpa
75.9/67
Schuchertinia conchicola (iso 132)
Schuchertinia_epiconcha_089 (short 28S)
100/100
Schuchertinia_epiconcha_112 (short 28S)
99.9/99
NH0434_Bonneviactinia_arctica
93.6/95
NOAH1154_Bonneviactinia_allmanii
97.2/100
69.4/73
NH0814_Bonneviactinia_allmanii
35/50
NOAH0621_Clava_multicornis
100/100
Clava multicornis (iso 139)
NH1049_Podocoryna_carnea
97.9/100
NH1030_Podocoryna_carnea
96.4/100
95.6/93
CC0316_Podocoryna_exigua
96.1/95
NH1357_Podocoryna_borealis
92.7/96
20.2/38
NH0134_Podocoryna_borealis
NH0129_Podocoryna_areolata
100/100
12.5/34
HYPNO_0595_Podocoryna_areolata
chimera_Bouillonactinia_hooperi (iso 205 + 238)
100/100
chimera_Bouillonactinia_carcinicola (iso 199 + H16S_1T7_
NOAH0705_Leuckartiara_nobilis
100/100
NOAH0135_Catablema
0.01
